# Structures of the Tilapia Lake Virus replication complex show that ANP32 is a host factor for replication across the *Articulavirales* order

**DOI:** 10.64898/2026.09.16.751959

**Authors:** Benoît Arragain, Martin Pelosse, Stephen Cusack

**Affiliations:** European Molecular Biology Laboratory, 71 Avenue des Martyrs, CS 90181, 38042 Grenoble Cedex 9, France; Institute of Science and Technology Austria, Am Campus 1, 3400 Klosterneuburg, Austria

## Abstract

Tilapia Lake Virus (TiLV) is a species in the Amnoonviridae family within the Articulavirales order of nuclear-replicating, segmented, negative-strand RNA viruses. This order also encompasses the well-known, but phylogenetically distant Orthomyxoviridae family that includes influenza and Thogoto viruses. Influenza virus replication, but not Thogoto virus, essentially relies on the host acidic nuclear phosphoprotein 32 (ANP32) to form a functional genome replication complex, which comprises an asymmetric viral polymerase dimer composed of a replicase bridged by ANP32 to an encapsidase. However, remains unclear whether this dependency is conserved in divergent viruses from the same order. Using in vitro reconstitution, biophysical and biochemical analyses and high-resolution structure determination by single-particle cryo-electron microscopy, we investigate the complexes formed between the TiLV RNA-dependent RNA polymerase (TiLV-Pol) and either tilapia (ti) or human (hu) ANP32A. We show that the tiANP32 leucine-rich repeat domain acts as a scaffold that stabilizes apo-TiLV-Pol in an encapsidase conformation. The tiANP32-encapsidase complex is then able to recruit and stabilise a second TiLV-Pol in a replicase conformation. The tiANP32-stabilised asymmetric TiLV-Pol dimer has an architecture that is remarkably similar to its orthomyxovirus replication complex counterpart, despite the 40% smaller size of TiLV-Pol. We also demonstrate that the ANP32 low-complexity acidic region interacts with the TiLV nucleoprotein, consistent with a conserved role in co-ordinating ribonucleoprotein assembly. Overall, our findings suggest that ANP32 is an ancient and likely broadly conserved host factor for viral genome replication and encapsidation across the Articulavirales order. Furthermore, TiLV is an emerging fish pathogen causing high mortality in aquaculture and our results provide insight into tilapia ANP32 modifications that could be used to engineer TiLV-resistant tilapia.

**AUTHOR SUMMARY:** The *Articulavirales* order of nuclear-replicating, segmented, negative-strand RNA viruses includes both the well-known Orthomyxoviruses such as influenza and Thogoto viruses and the recently discovered and distantly related *Amnoonviridae* family of mainly fish viruses, such as Tilapia Lake Virus (TiLV). It is thought that most orthomyxoviruses depend on host factor acidic nuclear phosphoprotein 32 (ANP32) to form the active genome replication complex. This comprises an asymmetric dimer of two viral polymerases (a replicase and an encapsidase) bridged by ANP32. Using *in vitro* biochemical, biophysical and high-resolution structure-determination methods, we investigate whether TiLV also requires host ANP32 for replication. We find that TiLV polymerase (TiLV-Pol) interacts with tilapia ANP32A (tiANP32A) to form an asymmetric dimer that architecturally resembles the influenza virus replication complex, despite TiLV-Pol being only 60% of the size of influenza polymerase and completely diverged in sequence. Furthermore, the C-terminal acidic tail of ANP32 binds TiLV nucleoprotein, suggesting that ANP32 could also be important for progeny genome encapsidation. These finding suggests that ANP32 is an ancient and likely broadly conserved host factor for viral replication across the *Articulavirales* order. Our results also show how genetic engineering of fish ANP32 could potentially be used to combat economically damaging fish diseases.

## INTRODUCTION

Tilapia Lake Virus (TiLV) is a recently discovered fish pathogen associated with high mortality rates in both wild and farmed tilapia^1,2^. TiLV is classified in the *Amnoonviridae* family, within the *Articulavirales* order of segmented negative-stranded RNA viruses (sNSV)^3,4^. This order also includes the *Orthomyxoviridae* family, which contains important human and animal pathogens such as influenza viruses, Thogoto virus (THOV), and infectious salmon anaemia virus (ISAV). Despite extreme phylogenetic divergence from classical orthomyxoviruses, the TiLV genome organisation and replication machinery share several conserved characteristics. The TiLV genome is divided into 10 single-stranded RNA segments, each with conserved, quasi-complementary 5′ and 3′ ends^3,5^. These segments encode putative viral proteins, most of which lack homology to known proteins, except for segment 1, which contains the conserved motifs characteristic of an orthomyxovirus PB1-like polymerase core subunit^3,5^.

Bioinformatic and structural analyses show that the TiLV RNA-dependent RNA polymerase (TiLV-Pol) is a heterotrimer comprising proteins encoded by segments 1-3^5^, while the viral nucleoprotein (TiLV-NP) is encoded on segment 4^6,7^. Despite their smaller size and sequence divergence, TiLV-Pol and TiLV-NP share structural similarities with their influenza counterparts, containing minimised versions of most corresponding domains^5,7^. TiLV-Pol binds the conserved 5′ and 3′ ends of the viral or complementary RNA (vRNA or cRNA) in a similar manner to the influenza viral promoter, with a 5′ hook and distal duplex region, in either mode A (with the 3′ end in the polymerase active site) or mode B (with the 3′ end in the secondary binding site)^5^. TiLV-Pol is active for RNA synthesis *in vitro* and can adopt distinct “transcriptase” and “replicase” conformations^5^. TiLV-NP can form various small, pseudo-symmetrical ring-like oligomers in the presence of RNA *in vitro*^7^, which have been structurally characterized using cryo-EM. This revealed the complete TiLV-NP structure, its oligomerization mode, as well as the RNA binding mode and directionality^7^, which are critical to the understanding of viral genome encapsidation into ribonucleoprotein particles (RNPs).

RNPs are the functional templates for both transcription and replication of the viral genome, two distinct RNA synthesis mechanisms that, in the case of influenza viruses, essentially depend on different host factors. Transcription relies on the direct interaction of the influenza polymerase (FluPol) with the host RNA polymerase II^8–10^, enabling cap-snatching of capped primers for transcription initiation. However, for TiLV-Pol, the PA endonuclease-like domain (PA/ENDO) is biochemically inactive and the PB2 cap-binding like domain (PB2/CBD) unable to bind m^7^GTP *in vitro*, raising doubts as to whether TiLV performs cap-snatching^5^. Replication by FluPol requires the acidic nuclear protein 32 (ANP32)^11^. ANP32 proteins are highly conserved amongst eukaryotes and perform diverse cellular functions as « electrostatic chaperones », notably of histones^12^. They comprise a folded N-terminal leucine-rich repeat (LRR) domain and a C-terminal intrinsically disordered low complexity acidic region (LCAR). The two distinct domains of ANP32 play crucial roles in the influenza viral replication process^11–13^. The LCAR is thought to first chaperone newly synthesised apo-FluPol, stabilising the encapsidase conformation, while the LRR bridges it to a promoter-bound FluPol within an RNP, forming an asymmetric dimer known as the replication complex. In this complex, the promoter-bound FluPol adopts the replicase conformation and synthesises a copy of the viral genome (either v- or cRNA), while the encapsidase is thought to bind the emerging 5′ end of the nascent RNA product. The LCAR domain is also proposed to recruit NPs onto the nascent RNA product that bulges out of the replication complex, progressively forming a progeny RNP^16^. To date, the *in vitro* reconstituted replication complex structures of influenza A^14,15^, B^15^, and C^13^ viruses have been described bound to human (hu)ANP32A, huANP32B or chicken (ch)ANP32A. These structures provide new insights into the influenza replication mechanism and rationalise the various adaptive mutations that avian FluPol can acquire to overcome host restriction. However, it is unclear whether ANP32 proteins are conserved host factors across the *Articulavirales* order, even though recent studies have investigated their implication in the replication of various Thogotovirus species^17,18^.

Here, we use *in vitro* complex reconstitution and mass photometry to show that the ANP32 LRR domain is sufficient to induce TiLV-Pol dimerization. Using single-particle cryo-electron microscopy, we then determined multiple high-resolution structures of monomeric or dimeric TiLV-Pol bound to either tilapia or human ANP32A (tiANP32A; huANP32A). We show that the ANP32 LRR domain acts as a scaffold that stabilizes apo-TiLV-Pol in an encapsidase conformation. This ANP32-encapsidase heterodimer is then able to recruit a second TiLV-Pol in the replicase conformation to form a ternary replication complex, an asymmetric dimer bridged by ANP32, whose architecture is remarkably similar to its influenza counterpart, despite TiLV-Pol′s minimal size and complete sequence divergence. We solve the TiLV replication complex structure in both apo and promoter-bound forms, and structurally map the interfaces that drive its assembly. Based on these high-resolution structures, we designed mutations in tiANP32A to disrupt TiLV-Pol-tiANP32A interactions *in vitro*, providing a structural basis for generating genetically-modified, disease-resistant tilapia. Finally, we show that the LCAR domain of tiANP32A is able to interact with TiLV-NP *in vitro*, supporting its putative role in NP recruitment during replication^16^. Overall, these results highlight that this highly conserved eukaryotic host factor was likely recruited very early in the evolution of articulaviruses, and articulaviral polymerases have evolved to maintain it as an essential component of the viral genome replication machinery.

## RESULTS

### The ANP32A LRR domain interacts with and mediates TiLV polymerase dimerization *in vitro*

To determine whether ANP32A can interact with TiLV-Pol and possibly induce dimerization, we first employed mass photometry (Fig. 1). In the presence of a 40-mer vRNA loop promoter (comprising the first and last 20 nucleotides of the 5′ and 3′ ends of TiLV segment 9^5^), TiLV-Pol is found to be monomeric with a measured mass of 157 kDa (Fig. 1A), consistent with previous single-particle cryo-EM experiments^5^. However, in the presence of either tilapia or human ANP32A, while the monomeric population remains, a second, lower-abundance population is observed that would correspond to a dimer of TiLV-Pol bridged by ANP32A (∼339-358 kDa) (Fig. 1B, C).

**Fig. 1.**
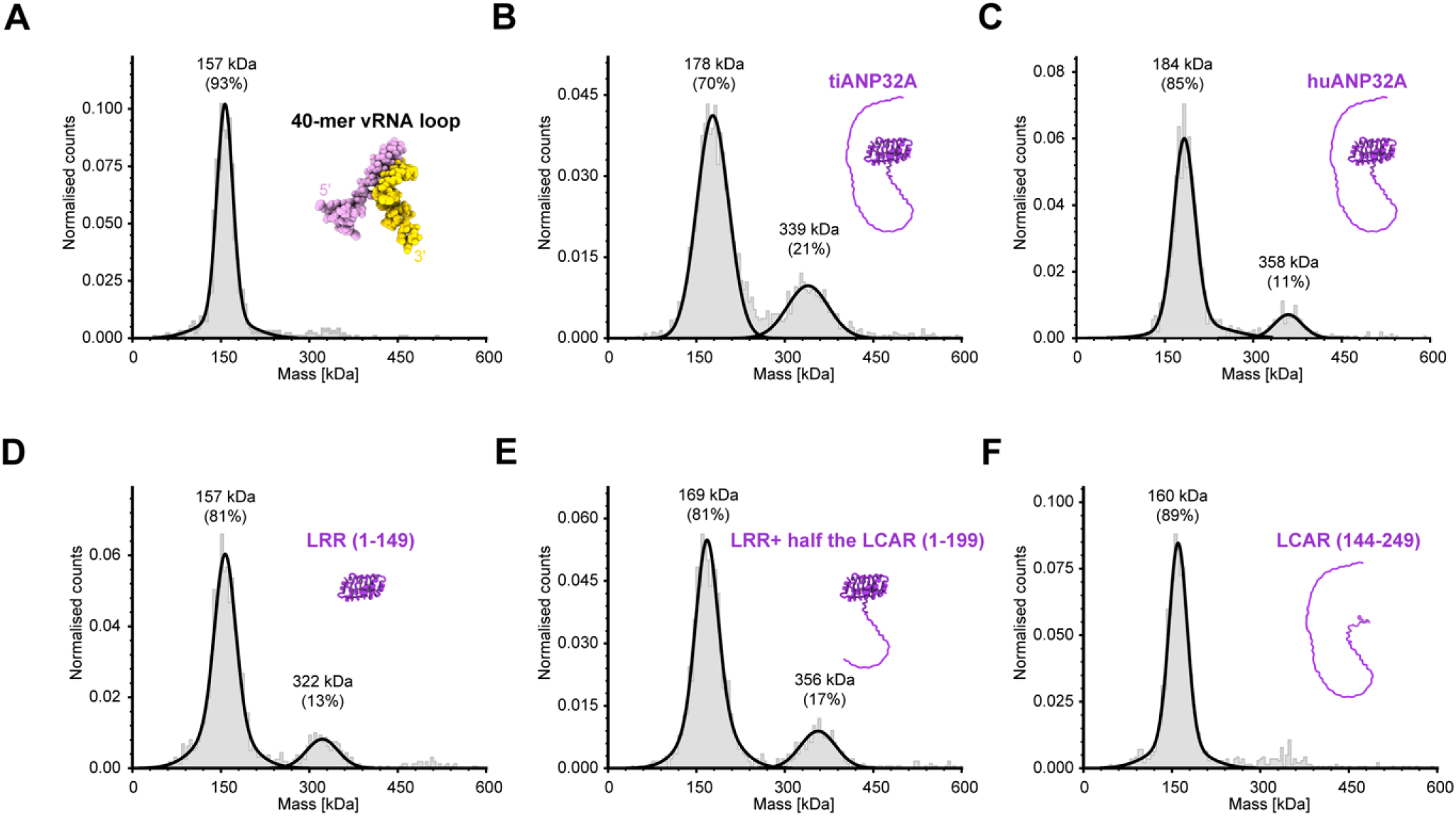
The ANP32A LRR domain interacts with and induces TiLV polymerase dimerization *in vitro*. Mass photometry analysis of TiLV-Pol mixed with (**A**) a 40-mer vRNA loop, (**B**) tilapia (ti)ANP32A, (**C**) human (hu)ANP32A, (**D**) the huANP32A leucine-rich repeat (LRR) domain (residues 1-149), (**E**) the huANP32A LRR domain plus half of the low-complexity acidic region (LCAR) (residues 1-199), and (**F**) the huANP32A LCAR alone (residues 144-249). Plots display normalized counts versus mass (kDa). Solid black lines indicate the main peaks. The estimated mass and relative percentage for each population are annotated.

To further identify the ANP32 domains responsible for TiLV-Pol dimerization, we tested complex formation using different truncated constructs of huANP32A. When using the LRR domain alone (1-149) or the LRR domain plus half of the LCAR (1-199), the TiLV-Pol dimeric population persisted (Fig. 1D, E). In contrast, using the LCAR alone (144-249) was insufficient to induce detectable TiLV-Pol dimerization (Fig. 1F). Taken together, these results indicate that TiLV-Pol dimerization is ANP32A-dependent and driven by the LRR domain.

Given the 91%/98% sequence identity/similarity between the huANP32A and tiANP32A LRR domains (Fig. S1), we anticipated a conserved interaction for both huANP32 and tiANP32A. We validated this by solving multiple single-particle cryo-EM structures at overall resolutions ranging from 2.4 to 3.2 Å of TiLV-Pol in complex with either ANP32A variant (summarized in Table 1; Fig. S2-S8; Tables S1-3). Two types of complex were obtained: (i) a monomeric TiLV-Pol bound to the ANP32A LRR, in which the host factor stabilizes the polymerase in an encapsidase conformation; and (ii) an asymmetric TiLV-Pol dimer bridged by ANP32A, solved in both apo- and vRNA promoter-bound states. The architecture of both complexes is highly reminiscent of previously reported ANP32-bound influenza polymerase complexes.

**Table 1.**
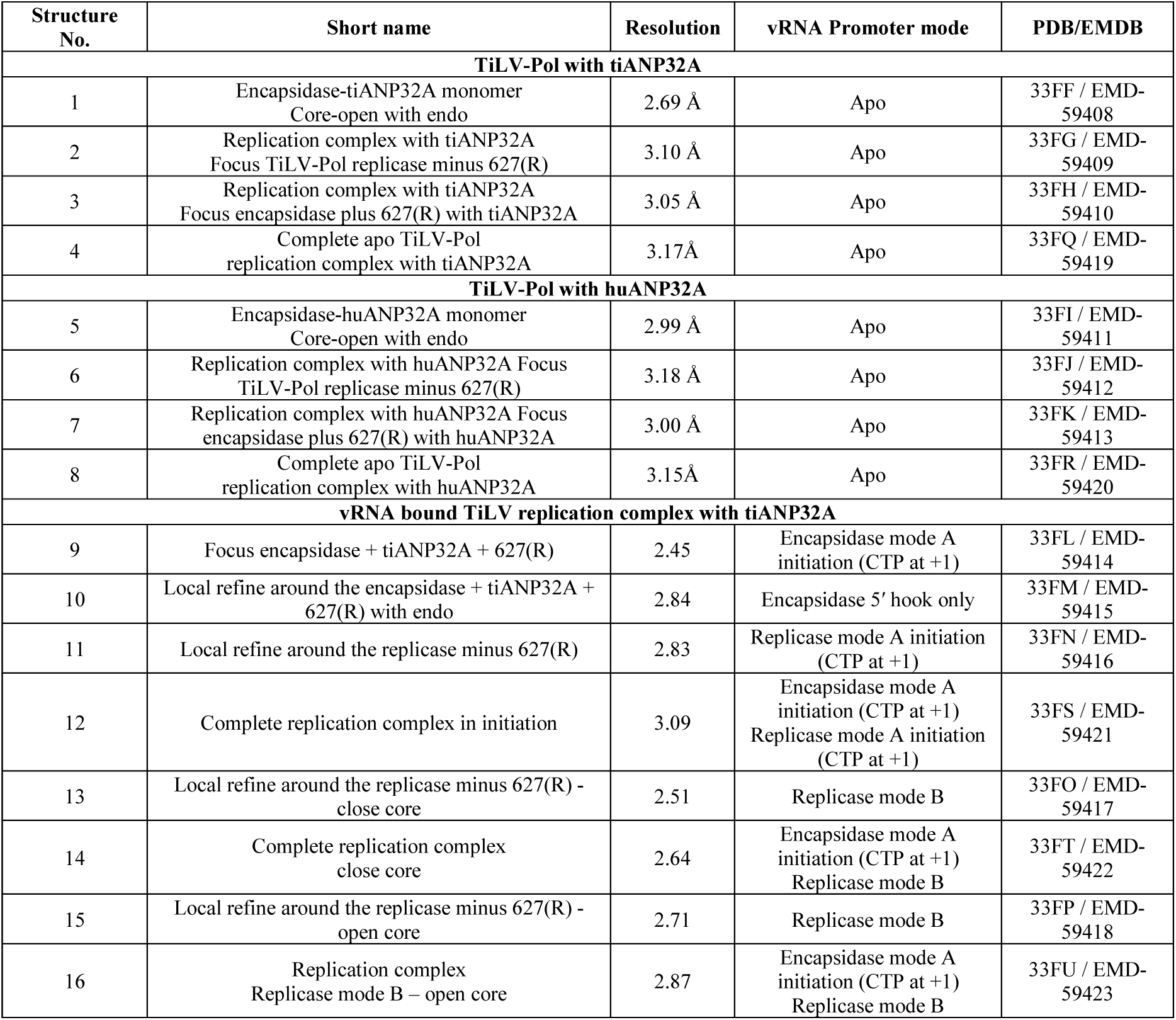
Summary of TiLV polymerase structures with tiANP32A or huANP32A.

| Structure No. | Short name | Resolution | vRNA Promoter mode | PDB/EMDB |
| --- | --- | --- | --- | --- |
| **TiLV-Pol with tiANP32A** |  |  |  |  |
| 1 | Encapsidase-tiANP32A monomer<br>Core-open with endo | 2.69 Å | Apo | 33FF / EMD-59408 |
| 2 | Replication complex with tiANP32A<br>Focus TiLV-Pol replicase minus 627(R) | 3.10 Å | Apo | 33FG / EMD-59409 |
| 3 | Replication complex with tiANP32A<br>Focus encapsidase plus 627(R) with tiANP32A | 3.05 Å | Apo | 33FH / EMD-59410 |
| 4 | Complete apo TiLV-Pol<br>replication complex with tiANP32A | 3.17Å | Apo | 33FQ / EMD-59419 |
| **TiLV-Pol with huANP32A** |  |  |  |  |
| 5 | Encapsidase-huANP32A monomer<br>Core-open with endo | 2.99 Å | Apo | 33FI / EMD-59411 |
| 6 | Replication complex with huANP32A Focus<br>TiLV-Pol replicase minus 627(R) | 3.18 Å | Apo | 33FJ / EMD-59412 |
| 7 | Replication complex with huANP32A Focus<br>encapsidase plus 627(R) with huANP32A | 3.00 Å | Apo | 33FK / EMD-59413 |
| 8 | Complete apo TiLV-Pol<br>replication complex with huANP32A | 3.15Å | Apo | 33FR / EMD-59420 |
| **vRNA bound TiLV replication complex with tiANP32A** |  |  |  |  |
| 9 | Focus encapsidase + tiANP32A + 627(R) | 2.45 | Encapsidase mode A<br>initiation (CTP at +1) | 33FL / EMD-59414 |
| 10 | Local refine around the encapsidase + tiANP32A +<br>627(R) with endo | 2.84 | Encapsidase 5' hook only | 33FM / EMD-59415 |
| 11 | Local refine around the replicase minus 627(R) | 2.83 | Replicase mode A initiation<br>(CTP at +1) | 33FN / EMD-59416 |
| 12 | Complete replication complex in initiation | 3.09 | Encapsidase mode A<br>initiation (CTP at +1)<br>Replicase mode A initiation<br>(CTP at +1) | 33FS / EMD-59421 |
| 13 | Local refine around the replicase minus 627(R) -<br>close core | 2.51 | Replicase mode B | 33FO / EMD-59417 |
| 14 | Complete replication complex<br>close core | 2.64 | Encapsidase mode A<br>initiation (CTP at +1)<br>Replicase mode B | 33FT / EMD-59422 |
| 15 | Local refine around the replicase minus 627(R) -<br>open core | 2.71 | Replicase mode B | 33FP / EMD-59418 |
| 16 | Replication complex<br>Replicase mode B – open core | 2.87 | Encapsidase mode A<br>initiation (CTP at +1)<br>Replicase mode B | 33FU / EMD-59423 |

### ANP32A stabilises monomeric apo-TiLV polymerase in an encapsidase conformation

Following complex reconstitution *in vitro* by mixing excess of ANP32A with apo TiLV-Pol (see Material and Methods), we solved the structures of the monomeric apo TiLV-Pol encapsidase (TiLV-Pol(E)) in complex with either tiANP32A (Table 1; Structure 1) or huANP32A (Table 1; Structure 5) at global resolutions of 2.7 Å and 3 Å, respectively (Table 1; Fig. 2; Fig. S2-S5). The ANP32-bound apo TiLV-Pol(E) heterodimers are predominant over the monomeric TiLV-Pol particles without ANP32A (Fig. S2, S4). As expected from the very high sequence similarity of their LRR domains (Fig. S1), the binding mode of both hu/tiANP32A is almost identical with all residues involved in the interaction with TiLV-Pol(E) being conserved (Fig. S1, S9). The only noticeable difference lies in the orientation of the solvent-exposed part of the LRR domain, which shifts by ∼6 Å (RMSD < 2.5 Å), whilst the PB2/627-NLS(E) double domain remains similarly organized (RMSD < 0.5 Å) (Fig. S9). In those structures, the PB2/Mid-link(E) and PB2/CBD(E) domains (residues 144-271) are not tightly integrated into the complex and are poorly visible due to flexibility (Fig. 2).

**Fig. 2.**
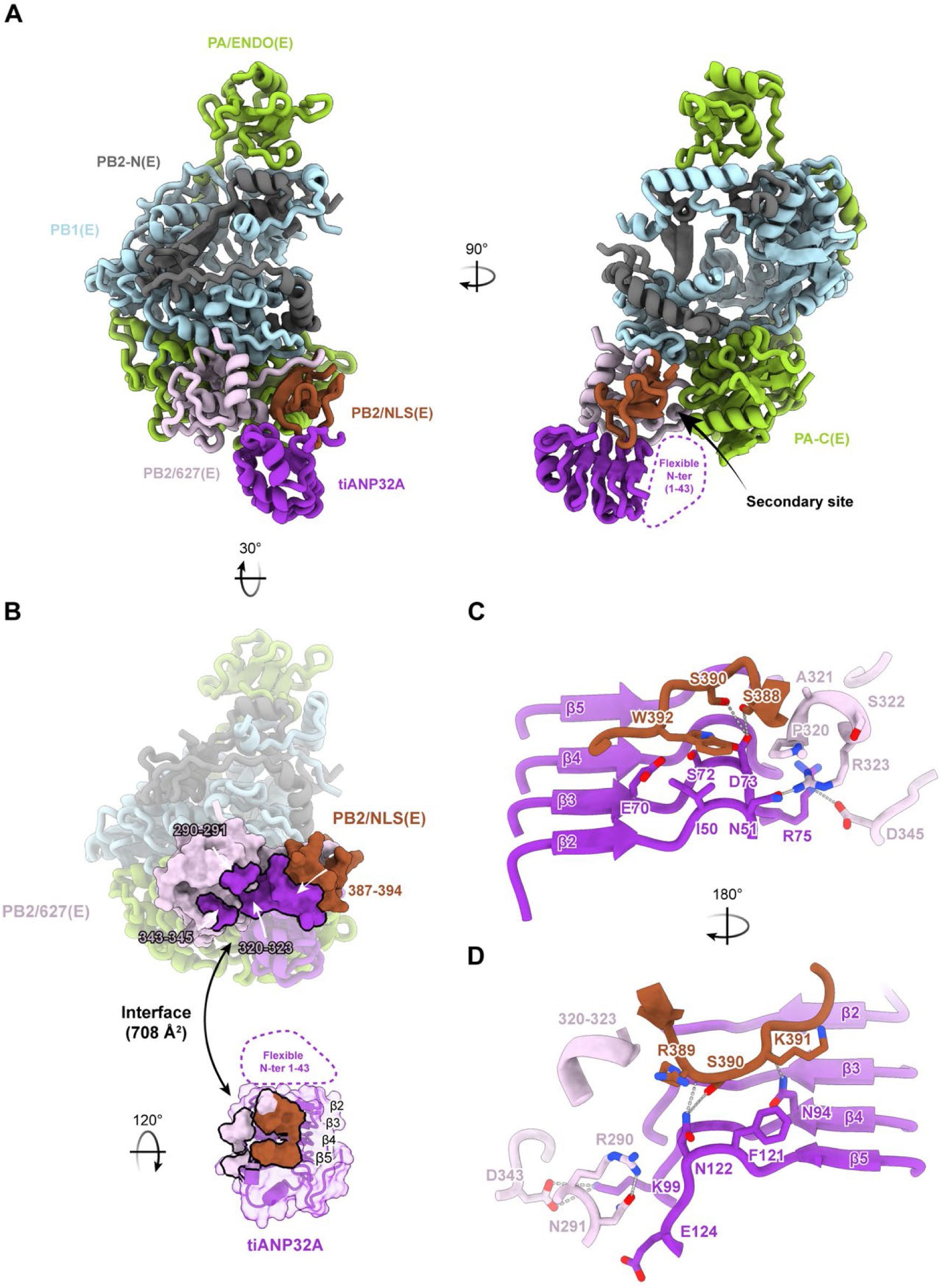
ANP32A stabilizes apo-TiLV polymerase in an encapsidase conformation. (**A**) Structure of the monomeric apo TiLV polymerase in encapsidase conformation bound to tilapia ANP32A with open core. The complex is shown in cartoon representation and coloured as follows: PA is light green, PB1 is light blue, PB2-N is dark grey, PB2/627 is light pink, PB2/NLS is brown, and tiANP32A is purple. The PB2/CBD domain, PB2/Mid-link domain, and tiANP32A N-terminus (residues 1-43) are not visible due to flexibility. The secondary RNA-binding site is indicated with an arrow. (**B**) View of the interface between TiLV-Pol(E) and tiANP32A. Top: the complex is rotated 30° relative to panel **A**, and the PB2/627-NLS(E) double domain is displayed as a surface with the tiANP32A footprint highlighted in purple. Bottom: the isolated tiANP32A is rotated 120°, with its interacting surface coloured according to the contacting domains on PB2/627-NLS(E). The total interface area is indicated. (**C-D**) Detailed molecular interactions between TiLV-Pol(E) and tiANP32A. Interacting residues are shown as sticks, and the tiANP32A β-strands are labelled. Hydrogen bonds and salt bridges are indicated by dotted lines.

In the ANP32-bound apo TiLV-Pol(E) heterodimers, the TiLV-Pol(E) conformation is stabilised by the interaction of the ANP32 LRR domain with the PB2/627-NLS(E) double domain (residues 276-422), which in turn docks onto PA-C(E) and PB1-C(E) (Fig. 2A). The interaction between the ANP32A LRR domain and TiLV-Pol(E) accounts for a total interface area of 883.7 Å^2^ (Fig. 2B). The N-terminal region of ANP32A is poorly resolved and makes minimal contact with PA-C(E), contributing only 20% (174.9 Å^2^) of this interface. In contrast, the central β-sheet region of the ANP32A LRR domain forms an extensive interface with the PB2/627-NLS(E) double domain, accounting for 80% (708.8 Å^2^) of the contact area (Fig. 2B-D). Specific interactions stabilising the complex starting from the end of the second LRR β-strand (LRR-β2), where I50 stacks against W392 of PB2/NLS(E). Nearby, N51 faces residues PB2/627(E) T319-P320 and forms a hydrogen bond with R323 (Fig. 2C). On LRR-β3, E70 and S72 also stack against PB2/NLS(E) W392, while D73 interacts with S388 and S390, and is sandwiched between PB2/627(E) P320 and PB2/NLS(E) W392. Additionally, R75 forms a hydrogen bond with PB2/627(E) D345 (Fig. 2C). On LRR-β4, N94 forms a hydrogen bond with the carboxyl group of PB2/NLS(E) K391, and K99 forms a salt bridge with PB2/627(E) D343 (Fig. 2D). Finally, on LRR-β5, F121 stacks against PB2/NLS(E) K391, N122 forms a hydrogen bond with the carboxyl group of PB2/NLS(E) R389 and S390 side chain, and E124 interacts with PB2/627(E) R290-N291 (Fig. 2D).

ANP32A-binding is coupled with the docking of the stabilized PB2/627-NLS(E) double domain onto the TiLV-Pol(E) core, spanning an interface area of ∼1223 Å^2^ (479 Å^2^ (39%) with PA-C(E) and 744 Å^2^ (61%) with PB1-C(E)) (Fig. S10). Specifically, PA-C(E) 314-317/342-347 residues interact with PB2/627(E) 352-360, while PA-C(E) E170/N246 interact with PB2/627(E) R317. Finally, PB1-C(E) residues 382-415 interact with the PB2/627-NLS(E) 362-381, C420, and K483 residues (Fig. S10). This overall structural arrangement effectively blocks the secondary binding site - which is normally occupied in pre-initiation and late-elongation states by the 3′ v/cRNA ends^5^ - due to slight rearrangement of specific PA-C(E), PB1(E), and PB1(E) residues (Fig. S10).

The identification of this ANP32-bound encapsidase conformation completes the repertoire of major functional states for TiLV-Pol, which depend on the structural flexibility of the PB2 C-terminal domains.

### All three functional polymerase conformations are conserved within the *Articulavirales* order

The primary feature that distinguishes the encapsidase, replicase, and transcriptase conformations is the distinct spatial arrangement of their PB2 C-terminal domains (Fig. 2; Fig. S11). In TiLV-Pol, this conformational plasticity is induced by the PB2/627-NLS linker (residues 375-382), which undergoes different structural rearrangements. In the previously characterized TiLV-Pol(T), the linker is compressed, resulting in a face-to-face arrangement of the PA/ENDO and PB2/CBD coupled with packing against the TiLV-Pol core of the PB2/Mid-link-627-NLS domains (Fig. S11A, D). This architecture is homologous to the FluPol(T) and linked to cap-snatching mechanism^19^ (Fig. S11D). Conversely, in the complete TiLV-Pol replicase (R) (i.e., extracted from the TiLV replication complex), the linker extends to project the PB2/NLS(R) onto PA/ENDO(R) while the PB2/Mid-link(R) and PB2/CBD(R) pack against the polymerase core, a rearrangement also observed in FluPol(R)^20,13–15^ (Fig. S11B, E). Distinct from both, ANP32 binding induces the encapsidase conformation in which the PB2/627-NLS(E) linker forms an additional β-strand that incorporates directly into the PB2/NLS(E) β-sheet, locking the PB2/627-NLS(E) double domain onto PA-C(E) and PB1-C(E) (Fig. S11C, F). Overall, the TiLV-Pol(E) conformation resembles recently characterized FluPol(E) structures, in which the PB2/627-NLS(E) double domain similarly packs against PA-C(E) and PB1(E)^13,15^ (Fig. S11F).

However, FluPol(E) exhibits additional contacts between PB2/CBD(E) and PA/ENDO(E), as well as the packing of PB2/Mid-link(E) onto PB2-N(E), which are not observed in the TiLV case due to the smaller size of its polymerase core. Of note, the complete FluPol(E) conformation has only been visualized within apo-FluPolB dimers^15^, bound to 5′ cRNA^15^, or within fully assembled replication complexes^13–15^. So far, only one other monomeric huANP32B-bound FluPolA/H5N1(E) structure analogous to TiLV-Pol(E) has been described, but the PB2 C-terminal domains were unresolved^14^. When comparing the host-factor interfaces, the main contact points between FluPol(E) and the ANP32 LRR domain are restricted to the 128-130 loop and a portion of the LRR β-sheet, yielding a total interface area of 1055.8 Å^2^. While the absolute ANP32-binding footprint is smaller in the ANP32A-TiLV-Pol(E) complex (883.7 Å^2^), since TiLV-Pol is only 60% the size of FluPol, this interface accounts for a proportionally larger portion of the TiLV-Pol(E) surface. This suggests that despite extreme sequence divergence, the minimal TiLV-Pol has evolved to maximize its dependence on the ANP32 LRR domain to stabilize the encapsidase conformation. The transcriptase, replicase and encapsidase conformations, thus likely exist in other viruses within the *Amnoonviridae* family and across the wider *Articulavirales* order.

### The N-terminal region of ANP32A bridges TiLV encapsidase and replicase to form a replication complex

In addition to the monomeric apo TiLV-Pol(E)-ANP32 structures presented above, we also solved the structures of ANP32A-bound TiLV-Pol asymmetric dimers, which we refer to as the TiLV replication complex (Fig. 3). This complex consists of an ANP32A-bound TiLV-Pol(E) bridged to a second TiLV-Pol in a complete replicase conformation. The replicase features a stabilized PB2/627(R) domain, with its PB2/NLS(R) peptide (residues 423-431) packing against the PA/ENDO(R) domain (Fig. 3A-C; Fig. S11). Two such structures, in complex with either tiANP32A (Table 1; Structure 4) or huANP32A (Table 1; Structure 8), were resolved at 3.17 Å and 3.15 Å, respectively (Table 1; Fig. S2-S8). To overcome inter-polymerase flexibility, particle subtraction and local refinements focusing on the “TiLV-Pol(R) minus 627(R)” (Table 1; Structures 2 and 6) and “TiLV-Pol(E)-ANP32A plus 627(R)” (Table 1; Structures 3 and 7) moieties allowed for substantial improvements in map quality (Table 1; Fig. S2-S5).

**Fig. 3.**
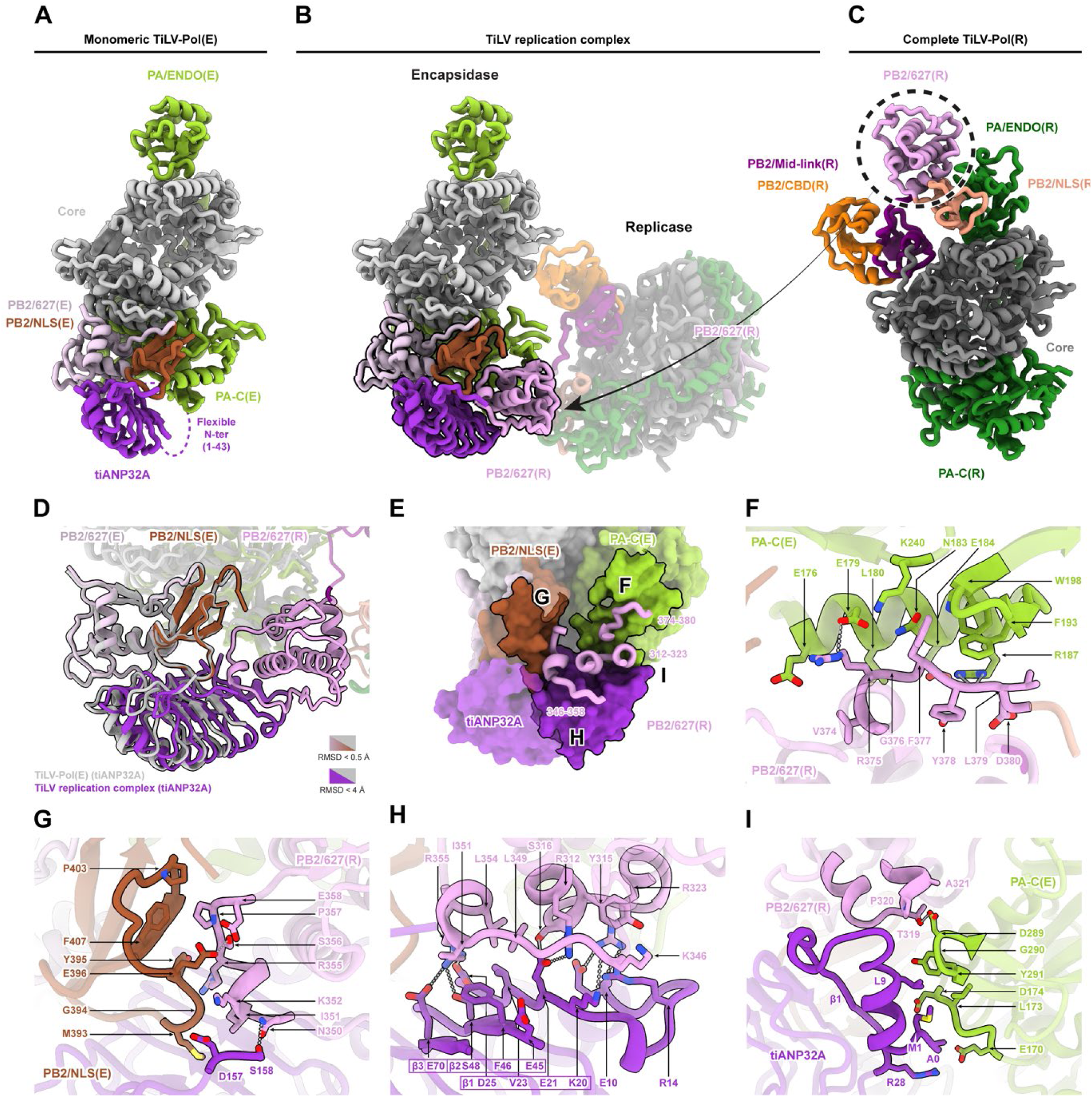
Architecture of the TiLV replication complex. (**A**) Structure of the monomeric apo TiLV polymerase in encapsidase conformation bound to tilapia ANP32A with open core. The complex is shown in cartoon representation and coloured as follows: PA is light green, the PB1 and PB2-N (core) is light grey, PB2/627 is light pink, PB2/NLS is brown, and tiANP32A is purple. The PB2/CBD domain, PB2/Mid-link domain, and the tiANP32A N-terminus (residues 1-43) are not visible due to flexibility. (**B**) Structure of the TiLV replication complex bound to tilapia ANP32A. The replicase moitie is shown as semi-transparent, with the exception of the PB2/627(R) domain. (**C**) Structure of the complete TiLV-Pol replicase extracted from the TiLV replication complex bound to tilapia ANP32A. TiLV-Pol(R) is shown in cartoon representation and coloured as follows: PA is dark green, the PB1 and PB2-N (core) is dark grey, PB2/627 is pink, PB2/NLS is beige, PB2/Mid-link is dark purple, and PB2/CBD is orange. The PB2/627(R) domain is surrounded by a dotted black circle, and a solid arrow indicates where it nests within the complete TiLV replication complex shown in panel **B**. (**D**) Superposition of the monomeric apo TiLV polymerase in encapsidase conformation bound to tilapia ANP32A onto the TiLV replication complex bound to tilapia ANP32A, aligned on the PB2/627-NLS(E) double domain. The monomeric encapsidase is coloured grey, while the replication complex is coloured as in panel **B**. The tripartite 627(E)-NLS(E)-627(R) assembly and the bridging tiANP32A are shown as non-transparent. The shift of tiANP32A upon 627(R) binding is visible (RMSD < 4.0 Å), while the 627-NLS(E) double domain organisation remains conserved (RMSD < 0.5 Å). (**E**) Close-up view of the PB2/627(R) interfaces with PA-C(E), PB2/NLS(E), and tiANP32A. The encapsidase domains and tiANP32A are shown as surfaces, while PB2/627(R) is shown as cartoon. Specific interacting regions corresponding to panels **F-I** are outlined in black. (**F-I**) Close-up views of the interfaces: (**F**) PA-C(E)-PB2/627(R), (**G**) PB2/NLS(E)-PB2/627(R)-tiANP32A, (**H**) PB2/627(R)-tiANP32A, and (**I**) PB2/627(R)-PA-C(E)-tiANP32A. Interacting residues are displayed as sticks and labelled. Hydrogen bonds and salt bridges are indicated by dotted lines.

In this asymmetric dimer, the TiLV-Pol(E) PB2/627-NLS(E) double domain maintains the exact same conformation observed in the monomeric encapsidase state (RMSD < 0.5 Å) (Fig. 3D). However, the previously poorly resolved N-terminus and central β-sheet of the ANP32A LRR domain is stabilised by a new interface of 493.8 Å^2^ with PB2/627(R). This allows PB2/627(R) to pack directly onto PB2/NLS(E) and PA-C(E), reinforcing the overall assembly through an extensive network of polar, hydrogen-bonding, and hydrophobic contacts across four distinct interaction hubs (Fig. 3E).

First, the PB2/627(R) loop (residues 374-380) makes extensive contact with PA-C(E) residues 176-198 (Fig. 3E, F). Hydrophobic contacts are formed between PB2/627(R) V374 and G376 and PA-C(E) L180. Nearby, PB2/627(R) R375 faces PA-C(E) E176 and forms a hydrogen bond with E179. PB2/627(R) F377 is locked in place: on one side by interactions with PA-C(E) N183, E184, and K240, and on the other by a cluster of aromatic residues (W198 and F193) (Fig. 3F). The aromatic ring of PB2/627(R) Y378 stacks in a cation-pi interaction against PA-C(E) R187, while its carboxyl group interacts with the same residue. Finally, PB2/627(R) L379 and D380 further lock this hub by interacting with PA-C(E) F193 and R187, respectively (Fig. 3F).

Second, on the opposite side, PB2/627(R) residues 350-358 interact with PB2/NLS(E) residues 393-407 and the C-terminal residues of tiANP32A, generating an interface area of 241.2 Å^2^ (Fig. 3G). These interactions involve PB2/627(R) residue N350, which interacts with the last visible C-terminal residue of the tiANP32A LRR domain S158. Concurrently, PB2/627(R) I351-K352 face PB2/NLS(E) M393 and tiANP32A D157. Finally, the PB2/627(R) 355-RSPE-358 motif engages directly with PB2/NLS(E) residues 394-GYE-396, as well as P403 and F407 (Fig. 3G).

Third, the anchoring of PB2/627(R) to PB2/NLS(E) and PA-C(E) positions PB2/627(R) residues 346-355 and 312-323 against the central β-sheet and N-terminus of tiANP32A (Fig. 3H). Starting from tiANP32A β3, residue E70 - in addition to its interaction with the monomeric TiLV-Pol(E) complex - forms a new hydrogen bond with PB2/627(R) R355. Residues on LRR β2 (E45, F46, and S48) and β1 (K20, E21, V23, and D25) face PB2/627(R) residues 346-355 and 312-323. Towards the tiANP32A N-terminus, E10 interacts with PB2/627(R) R323, which stacks with Y315. At the start of LRR β1, R14 and K20 face PB2/627(R) K346. E21 inserts into a pocket formed by S316 and L349, forming a salt bridge with R312. This interaction is strengthened via hydrophobic contacts between tiANP32A V23 and PB2/627(R) L354. Nearby, tiANP32A D25 forms a salt bridge with PB2/627(R) R355 within a cluster of polar residues that includes S48 (β2) and E70 (β3). Finally, tiANP32A E45 and F46 (β2) stack against PB2/627(R) I351 and L354 within a hydrophobic pocket (Fig. 3H).

Fourth and last, PB2/627(R) residues 319-321 and the N-terminal α-helix of the tiANP32A LRR domain (residues 0-28, with 0 corresponding to a leftover residue post-TEV cleavage) pack together onto PA-C(E) residues 170-174 and 288-291, effectively bridging the three domains (Fig. 3I). Specifically, PB2/627(R) T319 interacts with PA-C(E) D289, bringing G290 and Y291 into contact with the N-terminal α-helix of tiANP32A. In turn, PA-C(E) residues 170-174 face and interact with tiANP32A residues A0, M1, L9, and R28 (β1) (Fig. 3I).

Most reported interacting residues of tiANP32A LRR domain with TiLV-Pol(R) and TiLV-Pol(E) are conserved with other LRR domains from fish, human, or chicken, and if not, are of similar chemical properties (Fig. S1).

### Targeted tiANP32A mutations decrease TiLV replication complex formation *in vitro*

Based on our high-resolution structures, we investigated whether introducing mutations in tiANP32A could prevent the formation of the TiLV replication complex. We first designed various tiANP32A mutants targeting key residues within the salt-bridge interaction hubs at the interface with PB2/627(R) and PB2/NLS(E). To reduce Van der Waals interactions, we systematically introduced the F46A and F121A mutations. This double mutant (DM) was further combined with point mutations specifically designed to induce charge repulsion or steric clashes (e.g., DM combined with either E21K, D25K, I50W, E70K, or E73K).

Following expression and purification of the tiANP32A mutants, we conducted mass photometry experiments to assess whether these mutations abolish TiLV-Pol dimerization (i.e., replication complex formation). These experiments revealed reduced TiLV-Pol dimerization in the presence of all tiANP32A mutants *in vitro* (Fig. S12). Nevertheless, the extensive nature of the interface which spans the entire tiANP32A LRR domain suggests that achieving complete disruption may be challenging, and we cannot exclude that monomeric TiLV-Pol bound to tiANP32A mutants could remain in solution.

### The architecture of the TiLV replication complex is analogous to its orthomyxovirus counterparts

Compared to previously reported influenza replication complexes, the global organization of the TiLV replication complex is remarkably similar, sharing conserved structural features (Fig. S13).

First, the replicase displays a conserved interaction between the C-terminal PB2/NLS(R) α-helix and the PA/ENDO(R), a domain previously shown to be biochemically inactive *in vitro* in the case of TiLV-Pol^5^ and THOV-Pol^21^. Because this feature is absent in the structure of the monomeric (partial) TiLV-Pol(R) due to flexibility^5^, the formation of the full replication complex appears to stabilize this α-helical extension (Fig. S11). Second, ANP32 binding stabilizes the TiLV-Pol(E) conformation through interactions with the PB2/627-NLS(E) double domain, which in turn positions PB2/627(R) in close proximity, effectively sealing this tripartite assembly onto PA-C(E). Both the ENDO-NLS interaction and the tripartite assembly upon ANP32-binding represent hallmark features shared across the TiLV, FluA, FluB, FluC, and (to some extent) THOV replication complexes, despite phylogenetic divergence (Fig. S13).

Nevertheless, differences exist in the ANP32-polymerase interaction, which are primarily driven by the reduced size and sequence divergence of TiLV-Pol. In the TiLV replication complex, the ANP32A LRR domain interacts extensively from its N-terminus through to residue S158. In contrast, in the influenza A and B replication complexes, the main contact points between FluPol(E) and the ANP32 LRR domain - which rotates by ∼90° compared to TiLV - are restricted to the C-terminal half, which nests at the interface of the PA-C(E) 550-loop, PB1-N(E), PB2-NLS(E), and PB2/627(R)^14,15^, while for FluC, the ANP32 binding mode differs as well due to an altered PB2/627(R) orientation^13^ (Fig. S13).

Furthermore, while the tiANP32A LCAR points toward the TiLV PB2/627(R) domain, the density becomes unresolved beyond residue 158 and no additional density is observed elsewhere within the TiLV replication complex. This contrasts with the FluB replication complex, where the LCAR domain extends over the secondary binding site near the 5′ end, a trajectory supported by unique FluPolB PB1-C(E) and PB2-N(E) domains interacting with PB2/CBD(R)^15^.

These insights also contextualize the recently solved structure of an asymmetric dimer of THOV-Pol^17^. While closely resembling an influenza-like replication complex in its organization with similar replicase and encapsidase-like conformations and position, it assembled *in vitro* in the absence of ANP32, leaving the PB2/627-NLS(E) double domain flexible and unresolved (Fig. S13). This suggests that THOV replication could be ANP32 independent, consistent with the only 5-fold reduction in replication of

THOV in triple knockout huANP32A, B and E^17^ or double knockout huANP32A and B^22^ cells. Similarly, Dhori virus (DHOV), another Thogotovirus species, only exhibits a 10-fold reduction in replication in the absence of ANP32^22^. In contrast, Bourbon virus (BRBV), a relative of DHOV, also infecting both ticks and humans, absolutely requires huANP32A or B, or tick ANP32A, to replicate^22^. This is similar to FluA, FluB and FluC, even though FluC can form an asymmetric replication complex *in vitro* in the absence of any ANP32^13^.

To conclude, this distinct, yet structurally conserved replication complex formation and architecture - driven by ANP32 binding and stabilization of the encapsidase conformation - likely extends to other viruses within the *Amnoonviridae* family and the wider *Articulavirales* order, albeit with possible exceptions. It is also tempting to speculate on the formation of functionally analogous asymmetric dimers involved in replication within the distantly related *Bunyavirales* order as reported with Hantaan virus L protein^23^, though it remains to be determined whether host factors are similarly required to orchestrate their assembly.

### vRNA-binding by the replication complex, the product RNA trajectory, and LCAR-mediated nucleoprotein recruitment

To investigate whether the replicase and encapsidase components of the replication complex could bind vRNA promoters, we incubated TiLV-Pol with tiANP32A, a 40-mer vRNA loop (comprising the first and last 20 nucleotides of the 5′ and 3′ ends of TiLV segment 9), and CTP. Particle subtraction and local refinements focusing on the “TiLV-Pol(R) minus 627(R)” and “TiLV-Pol(E)-ANP32A plus 627(R)” subcomplexes, followed by 3D classification, allowed us to solve multiple TiLV-Pol replicase and encapsidase structures in different vRNA-binding modes at overall resolutions ranging from 2.5 to 2.9 Å (Table 1; Fig. 4; Fig. S6-S8).

**Fig. 4.**
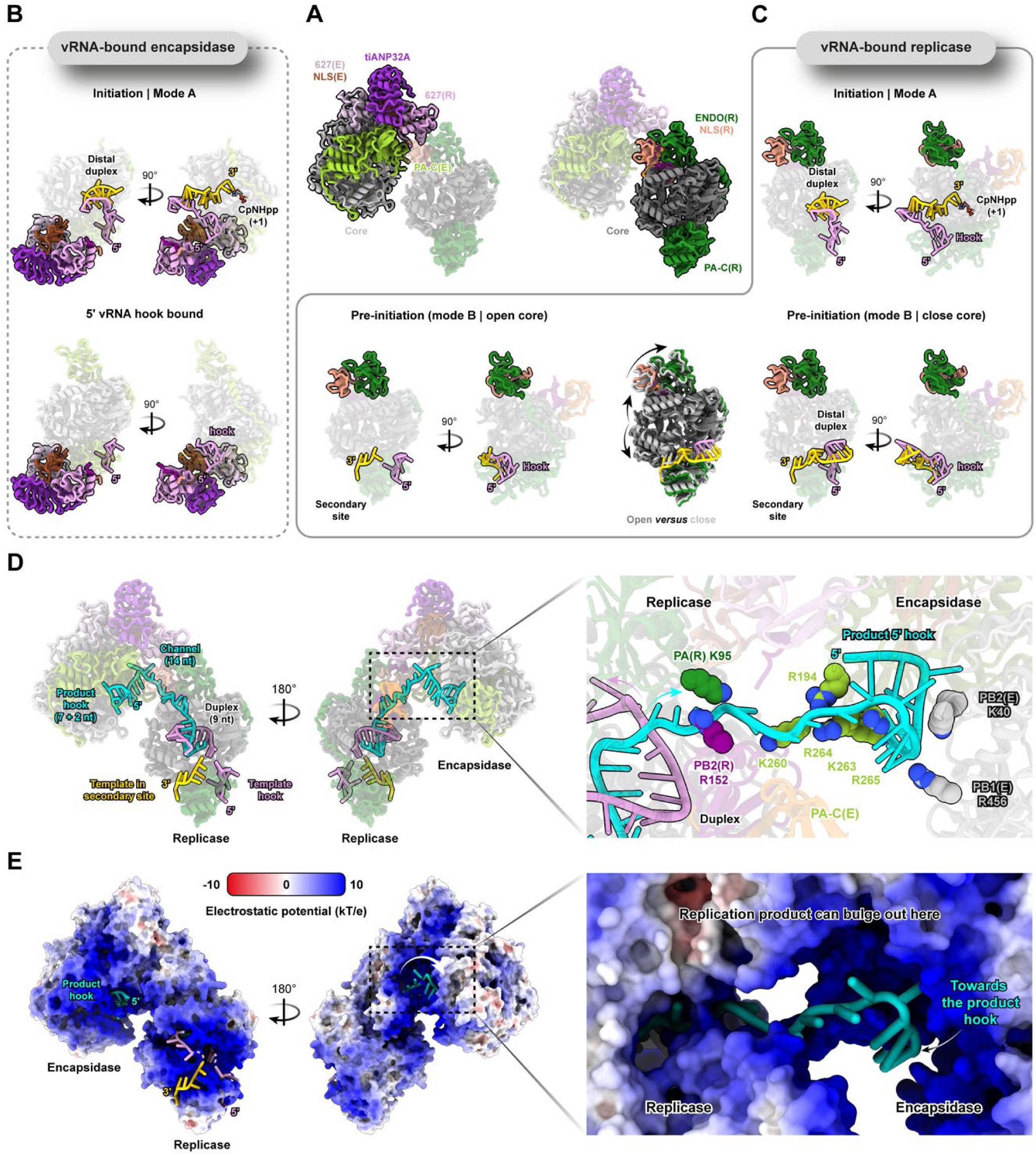
Structures of the vRNA-bound TiLV replication complex bound to tilapia ANP32A and model of an active elongation state. (**A**) Overview of the TiLV replication complex bound to tilapia ANP32A. The respective sub-complexes subjected to 3D classification and local refinements are shown as non-transparent cartoon: the encapsidase-tiANP32A plus 627(R) (left), and the replicase minus 627(R) (right). The encapsidase core is coloured light grey with PA-C(E) in light green, 627(E) in light pink, and NLS(E) in brown. The replicase core is dark grey, with PA/ENDO(R) in dark green, NLS(R) in beige, and 627(R) in pink. tiANP32A is purple. (**B**) Structures of the vRNA-bound encapsidase. Top: Initiation state (mode A) with both the 5′ and 3′ vRNA ends bound, forming a distal duplex, with CTP at the +1 active site position. Bottom: Encapsidase with only the 5′ vRNA hook bound. The 5′ vRNA is coloured plum and the 3′ vRNA is gold. (**C**) Structures of the vRNA-bound replicase. Top: Initiation state (mode A) with a distal duplex and CTP at the +1 position. Bottom left: Pre-initiation state (mode B) with an open core. Bottom right: Pre-initiation state (mode B) with a closed core and distal duplex formation. Bottom middle: Superposition of the open and closed core conformations. Black arrows highlight the core opening. (**D**) Cartoon representation of an active TiLV replication elongation complex, generated by superposing the open-core pre-initiation replicase, the pre-termination state (PDB 8QZ8), and the 5′ hook-bound encapsidase, illustrating the organization of the 5′ template hook, the 3′ template occupying the secondary site, the 9-nt RNA duplex within the replicase central cavity, and the nascent replication product reaching the encapsidase 5′ hook-binding site. The right panel provides a close-up view of the putative positively charged residues that guide the product towards the encapsidase. Residues are displayed as spheres and coloured by domain: PA(R) in dark green, PB2(R) in magenta, PA-C(E) in light green, and PB1/PB2(E) in light grey. (**E**) Electrostatic potential analysis of the modelled active TiLV replication elongation complex. The complex is shown in surface representation, coloured by electrostatic potential. The right panel shows a close-up view of the highly positive exit channel where the nascent RNA replication product (cyan) could bulge out between the replicase and encapsidase.

For the tiANP32A-TiLV-Pol(E)-627(R) subcomplex (Fig. 4A left, B), two different structures were obtained: Structure 9 in the initiation state (mode A) with CTP in the +1 position (Fig. 4B top), and Structure 10 where only the 5′ hook and the ENDO(E) is resolved (Fig. 4B bottom). For the TiLV-Pol(R) minus 627(R) moiety (Fig. 4A right, C), most particles were found in a pre-initiation state, with the 5′ hook bound and the 3′ end located in the secondary site (mode B), displaying varying degrees of core opening (Table 1; Structures 13 and 15; Fig. 4C). Two additional subpopulations were identified: TiLV-Pol(R) in pre-initiation mode B with a distal duplex (not refined); and Structure 11 an initiation state where the 3′ end is positioned in the PB1 active site with CTP at the +1 position (mode A) (Fig. 4C). In these vRNA-bound TiLV replication complex structures, the interactions between tiANP32A and TiLV-Pol(R/E), as well as those between TiLV-Pol(R/E) and the vRNA, remain strictly conserved relative to the apo-TiLV replication complexes and previously described TiLV-Pol-vRNA structures^5^. We note that in the different vRNA-bound replication complex structures, low resolution density is observed for the PB2/Mid-link(R) and PB2/CBD(R) domains (residues 144-271), but this flexibly linked domain does not appear to play any role in stabilising the replication complex.

Notably, no monomeric vRNA-bound TiLV-Pol(E) particles associated with ANP32A were observed in our datasets (Fig. S7). This suggests that TiLV-Pol(E) can only capture the 5′ and 3′ RNA ends when stabilized as part of the fully assembled replication complex. Furthermore, because the secondary binding site in TiLV-Pol(E) is sterically occluded by the PB2 C-terminal domains upon ANP32 binding, the 3′ end cannot access it and is instead projected directly into the PB1(E) active site. The observation that TiLV-Pol(R) and TiLV-Pol(E) can both have varying degrees of core opening and mode of promoter binding, shows that the overall replication complex architecture is robust towards the kinds of conformational changes that must occur during the different steps of replication.

By superposing the previously obtained TiLV-Pol pre-termination state (PDB 8QZ8) onto the open-core pre-initiation replication complex structure, and in combination with the 5′ hook-bound encapsidase structure, we modelled a putative, active TiLV replication complex with TiLV-Pol(R) bound to the promoter regions and accommodating a 9-bp RNA duplex within its central cavity (Fig. 4D, E). Extending the replication product RNA chain toward the 5′ hook binding site of TiLV-Pol(E) suggests that a minimum of ∼32 nucleotides would be required for the encapsidase to capture the newly synthesized 5′ end. This trajectory comprises the 9-bp duplex in the replicase cavity, 14 nucleotides spanning an open channel linking the replicase to the encapsidase, and 9 nucleotides corresponding to the bound hook (including the two unseen, TiLV specific 5′ terminal nucleotides^5^).

In this model, the product channel is lined with positively charged residues from both the replicase (PA(R) K95, PB2(R) R152) and the encapsidase (PA-C(E) K260, K263, R264, R265; PB2(E) K40; PB1(E) R456) that would primarily interact with the RNA phosphate backbone (Fig. 4E). Electrostatic potential analysis of the TiLV replication complex indeed reveals the presence of a highly basic open channel where the nascent replication product could bulge out during elongation, providing an accessible site where the ANP32 LCAR could also recruit NPs for genome encapsidation (Fig. 4D).

To link these structural observations to RNP genesis, we further investigated the putative interaction between ANP32A and the TiLV-NP. Having recently established expression and purification protocols for TiLV-NP^7^, we performed size-exclusion chromatography (SEC) experiments. Using wild-type tiANP32A and huANP32A, we observed a clear co-elution between TiLV-NP and both ANP32A variants (Fig. S14). We then utilized huANP32A truncated constructs (LRR alone 1-149; LCAR alone 144-249), which revealed that the TiLV-NP-ANP32A interaction is solely mediated by the LCAR domain, as no interaction was detected with the LRR domain alone (Fig. S14). In light of prior findings regarding the interaction between influenza A NP and ANP32 LCAR^16^, these results support the hypothesis that the intrinsically disordered LCAR may act as an “electrostatic whip” to recruit NP. By contacting the positively charged RNA-binding groove of apo-TiLV-NP, the LCAR could actively shuttle NPs onto the bulging RNA replication product, preventing non-specific RNA competition and promoting the processive encapsidation of the viral genome.

## DISCUSSION

In this study, we biochemically and structurally characterized the interactions between the conserved eukaryotic host factor ANP32 and the replication machinery of TiLV, a highly pathogenic member of the *Amnoonviridae* family that impacts tilapia aquaculture^2^. We demonstrate that despite extreme sequence divergence and minimal size compared to its orthomyxovirus counterparts, TiLV-Pol strictly depends on ANP32 for dimerization (Fig. 1). Specifically, the ANP32 LRR domain acts as a structural scaffold, interacting with and stabilizing apo-TiLV-Pol in an encapsidase conformation (Fig. 2). This apo-TiLV-Pol(E)-ANP32 subcomplex subsequently recruits a second TiLV-Pol (this would be RNP-resident in an infected cell) in a replicase conformation to assemble the functional asymmetric replication complex. Within this ANP32-bound asymmetric dimer, the replicase is characterised by the ENDO-NLS(R) interaction, together with the packing of the tripartite 627(R)/NLS(E)-627(E) assembly onto PA-C(E) (Fig. 3). The overall architecture of the TiLV replication complex, including these two structural features, is remarkably conserved compared to the previously reported influenza A, B, and C replication complexes (and, to some extent, THOV)^13–15,17^. Furthermore, the vRNA-bound replication complex structures allow us to model an elongation state, revealing a continuous, positively charged RNA path that could channel the nascent product from the replicase directly to the 5′ hook-binding site of the encapsidase (Fig. 4). Based on these structural insights and in the light of previous studies, we propose a general ANP32-dependent mechanism for Articulavirus genome replication (Fig. 5).

**Fig. 5.**
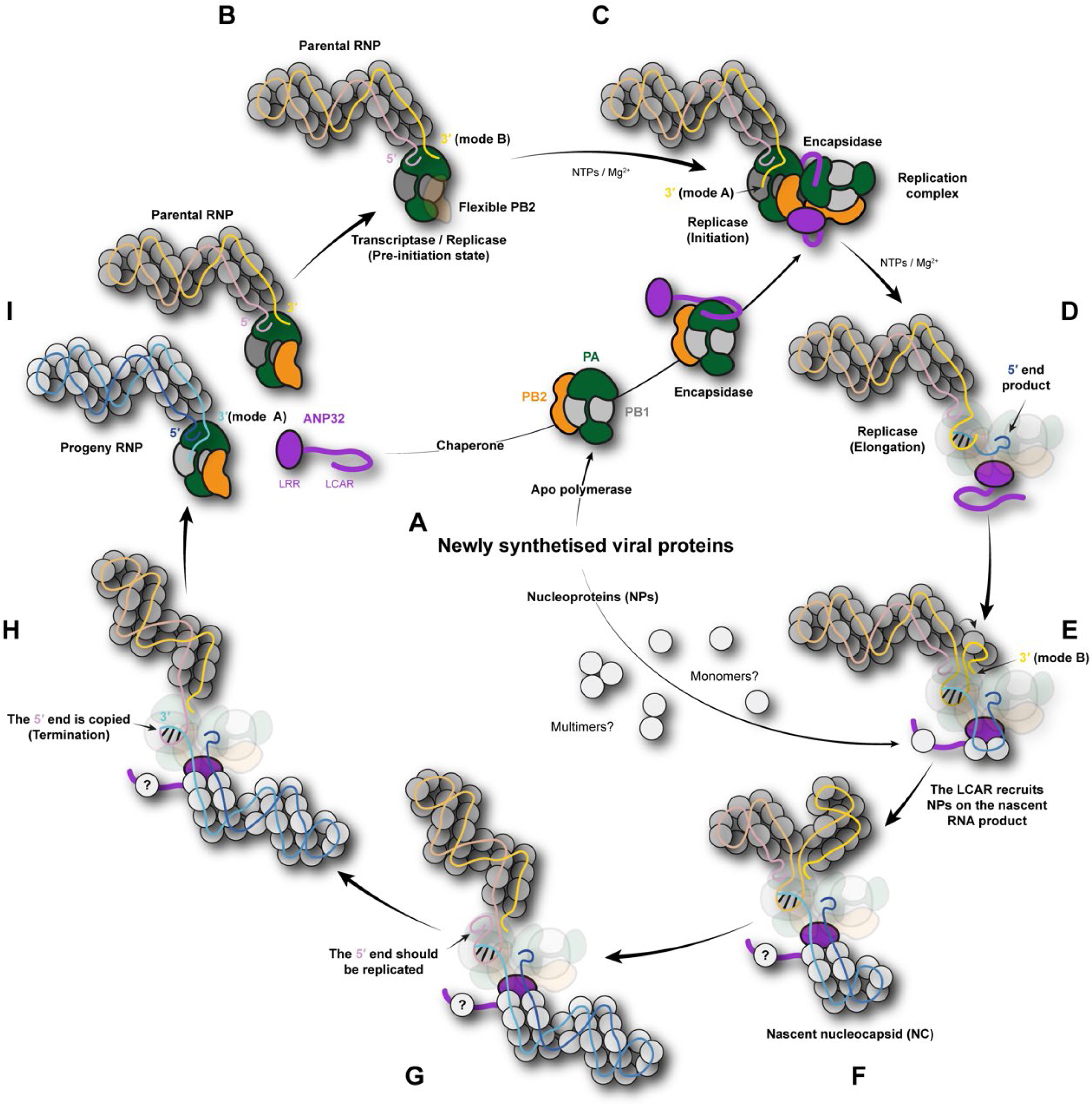
Viral genome replication mechanism of the *Amnoonviridae* and *Orthomyxoviridae* families. (**A**) Following viral mRNA transcription and translation, newly synthesized apo-polymerases are chaperoned by the host protein ANP32. Interactions with the ANP32 LRR and/or LCAR domains induce the apo-polymerase to adopt an encapsidase conformation. In the meantime, copies of the nucleoproteins (NPs) are also synthesized. (**B**) Within a parental RNP, following a transcription or replication cycle, a polymerase is bound to promoter regions with flexible PA-ENDO and PB2-C domains. (**C**) The ANP32-bound encapsidase docks onto the promoter-bound polymerase within a parental RNP, inducing the replicase conformation. This acts as a structural lock to stabilize the complete replication complex. Upon the influx of NTPs and Mg^2+^, the replicase enters the initiation state mode A. (**D**) The replicase proceeds to elongation. The nascent RNA product is extruded through a positively charged channel, guiding the 5′ end toward the hook-binding site of the encapsidase. (**E**) As elongation continues and the nascent RNA product bulges out, the ANP32 LCAR domain acts as an electrostatic whip to recruit apo-nucleoproteins (NPs) - which may exist as monomers or multimers - onto the nascent RNA. (**F-G**) The nascent nucleocapsid (NC) continues to form. It remains to be understood if the ANP32A LCAR acts as a NC nucleation point at the start of the replication and/or if the LCAR is needed all along the NC genesis (?). To complete genome replication, the replicase must copy right to the 5′ end of the template, necessitating the release of the tightly bound 5′ hook via a yet to be discovered mechanism. (**H**) During termination, the 5′ end is copied. The newly synthesized 3′ end of the product is projected directly toward the encapsidase active site, as the secondary site is inaccessible. (**I**) It is hypothesized that the engagement of the 3′ end with the encapsidase, coupled with polymerase core opening during elongation, would disrupt the asymmetric dimer. This triggers the release of ANP32 and separates the parental RNP from the progeny RNP, completing the replication cycle.

Following the transcription and translation of viral mRNAs, newly synthesized apo-polymerases are first chaperoned by ANP32^15^ (Fig. 5A). For FluPol, it was previously shown that the ANP32 LCAR is implicated in stabilizing apo-FluPol at physiological salt concentrations as symmetrical dimers (distinct for FluPolA and FluPolB), wherein in the case of FluPolB, one moiety is able to adopt a complete encapsidase conformation^15^. While no such symmetrical dimers were observed for TiLV-Pol *in vitro*, the ANP32 LRR domain alone is sufficient to bind and stabilize a complete monomeric TiLV-Pol encapsidase by locking the PB2/627-NLS(E) double domain onto PA-C(E), a hallmark feature also observed within encapsidases from influenza A, B, and C replication complexes. Current data suggest that ANP32 induces apo-polymerases to adopt an encapsidase conformation via its LRR and/or LCAR domains. Whilst promoter-bound polymerase within a parental RNP retains considerable flexibility of the ENDO and PB2-C domains (Fig. 5B), our structures demonstrate that an ANP32-bound encapsidase is able to successfully dock onto and stabilize a complete replicase conformation by primarily interacting with the PB2/627(R) and ENDO(R) domains (Fig. 5C).

In the complete TiLV replication complex, the N-terminus and central β-sheet of the ANP32 LRR integrate the PB2/627(R) into the assembly and the C-terminal PB2/NLS(R) α-helix packs directly onto the PA/ENDO(R), a structural feature conserved in all known orthomyxovirus replicase structures. A common functional role for this interaction might be to serve as a structural lock that stabilizes the replicase conformation, thereby favouring unprimed, *de novo* replication, regardless of polymerases ENDO activity (TiLV-Pol^5^ and THOV-Pol^21^ ENDO are biochemically inactive). However, for polymerases possessing an active ENDO, such as FluPol, this ENDO-NLS(R) interaction might fulfil an additional regulatory role by restricting ENDO flexibility and activity, preventing the non-specific cleavage of the nascent RNA replication product.

Following replication complex formation, and upon influx of NTPs and Mg^2^⁺ into the PB1 active site, the replicase enters the initiation state (mode A) (Table 1; Structure 12) and then proceeds to elongation (not structurally observed) (Fig. 5D). The nascent RNA product is presumably extruded through a positively-charged channel that guides the 5′ end toward the hook-binding site of the encapsidase. We estimate a minimum of ∼32 nucleotides are necessary to bridge this distance and bind the encapsidase, compared to the ∼36 nucleotides required for the bulkier FluPol^15^. As elongation continues and the RNA product begins to bulge out of the replication complex, the ANP32 LCAR domain could act as an electrostatic whip to recruit apo-NPs (which may exist as monomers or multimers, as observed for TiLV-NP *in vitro*)^7^ (Fig. 5E). It remains to be understood if the ANP32A LCAR acts as a nucleocapsid (NC) nucleation point at the start of the replication and/or if the LCAR is needed all along the NC genesis (Fig. 5F). To complete genome replication, the replicase must copy right to the 5′ end of the template, which requires release of the tightly bound 5′ hook (Fig. 5G). Exactly what stimulates this release during replication termination remains poorly understood for sNSV polymerases in general (Fig. 5H). The newly synthesized 3′ end of the product is sterically blocked from binding the secondary site of the encapsidase (mode B) in TiLV. Consequently, it is projected directly toward the PB1(E) active site. We hypothesize that once this 3′ end engages the encapsidase active site to initiate a new round of RNA synthesis, the resulting template-product duplex coupled with polymerase core opening in elongation, induces the disruption of the asymmetric dimer, releasing ANP32, the parental RNP, and the newly formed RNP (Fig. 5I). The encapsidase can then reconfigure into a transcriptase (after cRNA to vRNA synthesis) or a partial replicase for the next round of mRNA or genomic RNA synthesis.

The dependence of TiLV-Pol asymmetric dimerization on ANP32 highlights this protein as an anciently recruited host factor for viral genome replication. A recent phylogeny-based hypothesis propose that the *Articulavirales* order has an ancient aquatic origin^24^, with viral diversification linked to the emergence of aquatic animals. It is truly remarkable that over millions of years of evolution, Articulavirus polymerases appear to have maintained their dependence on ANP32 as an essential cofactor for replication. Specific viruses, for example for THOV^22,17^, may have idiosyncratically evolved away this dependence. In this case, stronger encapsidase-replicase interactions may be enough to stabilise the asymmetric dimer, but how the proposed recruitment of NP by the LCAR is bypassed is not clear. This conserved use of this host factor is all the more remarkable in that, whereas ANP32 has hardly diverged at all in size and sequence (e.g., >90% identity between vertebrate ANP32A, although invertebrate ANP32 sequences are more divergent), TiLV-Pol, in common with several other Amnoonviral polymerases, is considerably smaller than, for instance FluPol, yet has co-evolved to maintain the ability to functionally interact. Indeed the use of a highly conserved host factor perhaps facilitates cross species infection by certain Thogotoviruses that can infect both ticks and humans^17^, although host restriction by ANP32 can also exist, as for FluA between birds and mammals.

Whereas most classical orthomyxovirus polymerases are presumed to obtain capped primers for viral transcription by binding and cleaving nascent host Pol II transcripts, the Thogotoviruses as well as TiLV and perhaps other articulaviruses, exceptionally have biochemically defunct endonuclease and cap-binding domains^5,21,17^. Nevertheless, these domains have been maintained as they are required structurally to form the replication complex, especially the ENDO(R) domain (although in TiLV the CBD does not seem to be used for this). Interestingly, our structure of the TiLV replication initiation state, with incoming CTP at the +1 position, may give some insight into how capped mRNAs that lack 5′ end host sequences are generated in TiLV and possibly THOV. The vRNA configuration in the TiLV replication initiation state asymmetric dimer is identical to that in the monomeric TiLV transcription initiation state (PDB 8PSN), suggesting that in both cases RNA synthesis is terminally initiated on the vRNA template. We propose that in the case of the monomeric transcriptase, during early elongation the emerging 5′ end of the product can be capped by cellular capping enzymes (this might be facilitated by tethering the polymerase in the vicinity of the host Pol II^25^), thus producing a capped viral mRNA. In contrast, in the context of the full replication complex, the emerging 5′ end of the product is protected from solvent by channelling it into the encapsidase where the 5′ hook is tightly bound. This would be consistent with the lack of host sequences in THOV mRNAs^26^, although this has not been reported for TiLV yet. An alternative explanation for the lack of cap-snatching in these viruses would be that transcription is primed by host-produced capped dinucleotides^17^.

Our high-resolution structures also provide potential new avenues for TiLV disease control. Recent studies have demonstrated that CRISPR/Cas9 genome editing of ANP32A can confer partial resistance to avian influenza in poultry^27^. While genetically modifying chickens poses epidemiological risks, such as driving the emergence of virulent strains capable of cross-species transmission, TiLV is unlikely to ever be a risk to humans. Our demonstration that *in vitro* mutations of tiANP32A disrupt TiLV-Pol dimerization establishes a foundation for genetically engineering TiLV-resistant tilapia by modifying the tiANP32 gene(s), although care would be needed to maintain essential tiANP32 functions. Assuming its replication is also ANP32 dependent, a similar strategy may potentially be applicable to the salmon anaemia virus (ISAV), an orthomyxovirus pathogen of the Atlantic salmon that is a serious threat to the salmon farming industry^28^.

Future work will aim to capture structural snapshots of the TiLV replication complex in active elongation, and to validate our *in vitro* observations using cell-based minigenome systems, reverse genetics^29^ and ANP32-knockout fish cell lines^30^. These functional and virological assays will be critical to fully elucidate the dependence of the TiLV replication cycle on tiANP32, but are beyond the scope of the current *in vitro* study.

Finally, our findings provide compelling structural and biochemical evidence that the fundamental mechanisms of Articulavirus replication, from apo-polymerase stabilization to replication complex assembly and RNP encapsidation, are conserved.

## MATERIALS AND METHODS

### Cloning, expression and purification of the TiLV polymerase

TiLV polymerase (TiLV-Pol) (PB2 C-terminal deca-His-tag) was expressed and purified as previously described^5^. Briefly, for insect cells expression, *Trichoplusia ni* High 5 cells (ThermoFisher) at 0.8-1 × 10^6^ cells/mL concentration were infected by adding 1% of virus. The cells were disrupted by sonication in lysis buffer 50 mM HEPES pH 8, 500 mM NaCl, 20 mM imidazole, 0.5 mM TCEP and 10 % glycerol with cOmplete EDTA-free Protease Inhibitor Cocktail (Roche). After lysate centrifugation, the soluble fraction was loaded on a HisTrap HP ion affinity chromatography (Cytiva). Bound proteins were subjected to two sequential wash steps and eluted using initial lysis buffer supplemented by 500 mM imidazole. TiLV heterotrimeric complex fractions were pooled and diluted to reach the heparin-loading buffer concentration (50 mM HEPES pH 8, 250 mM NaCl, 0.5 mM TCEP, 5% glycerol). Proteins were loaded on a HiTrap Heparin HP (Cytiva) column, washed and eluted using 50 mM HEPES pH 8, 1 M NaCl, 2 mM TCEP, 5% glycerol. TiLV heterotrimeric complex fractions were then concentrated, flash frozen in liquid nitrogen, and stored at −80 °C for further use.

In addition, a new TiLV polymerase construct was cloned with a double strep-tag on PB2 C-terminal, instead of the deca-His-tag. For insect cells expression, *Trichoplusia ni* High 5 cells (ThermoFisher) at 0.8-1 × 10^6^ cells/mL concentration were infected by adding 1% of virus. The cells were disrupted by sonication in lysis buffer 50 mM HEPES pH 8, 500 mM NaCl, 2 mM TCEP and 10 % glycerol with cOmplete EDTA-free Protease Inhibitor Cocktail (Roche). After lysate centrifugation at 48,000 g for 45 min at 4 °C, the soluble fraction was loaded on strep-tactin affinity purification beads (IBA, Superflow). Bound proteins were eluted using the lysis buffer supplemented by 2.5 mM d-desthiobiotin (ThermoFisher). Protein-containing fractions were pooled, diluted to reach the heparin-loading buffer concentration (50 mM HEPES pH 8, 250 mM NaCl, 2 mM TCEP, 5% glycerol), and subsequently loaded on an affinity column HiTrap Heparin HP 5 mL (Cytiva). A continuous gradient from the heparin-loading buffer to the heparin-elution buffer (50 mM HEPES pH 8, 1 M NaCl, 2 mM TCEP, 5% glycerol) was applied over 15 column volume (CV). TiLV-Pol fractions were pooled and dialysed overnight in a final buffer (50 mM HEPES pH 8, 500 mM NaCl, 2 mM TCEP, 5% glycerol), concentrated with Amicon Ultra-15 (30 kDa cutoff), flash-frozen and stored at −80 °C for later use.

### Cloning, expression and purification of the TiLV nucleoprotein

TiLV nucleoprotein (TiLV-NP) (N-terminal deca-His-tag) was expressed and purified as previously described^7^. For large-scale expression, *Trichoplusia ni* High 5 cells (ThermoFisher) at a concentration of 0.8-1 × 10^6^ cells/mL were infected by adding 1% of the virus. Expression was stopped 72 to 96 hours after the day of proliferation arrest, and the cells were harvested by centrifugation (1000 g, 20 min, 4 °C). The cells were disrupted by sonication for 3 minutes (10 s ON, 20 s OFF, 40% amplitude) on ice in lysis buffer (50 mM HEPES pH 8, 500 mM NaCl, 20 mM imidazole, 0.5 mM TCEP, and 5% glycerol) with cOmplete EDTA-free Protease Inhibitor Cocktail (Roche). After lysate centrifugation at 48,000 g for 45 minutes at 4 °C, the soluble fraction was loaded on a HisTrap HP ion affinity chromatography column (Cytiva). Bound proteins were subjected to two sequential wash steps using (i) the lysis buffer supplemented with 1 M NaCl and (ii) the lysis buffer supplemented with 50 mM imidazole. Bound proteins were eluted gradually from 50 mM imidazole to 500 mM imidazole over 15 CV. TiLV nucleoprotein fractions were pooled. Tobacco Etch Virus protease was added for His-tag cleavage (1:50 w/w ratio), and the protein mixture was dialyzed overnight at 4 °C in a heparin-loading buffer (50 mM HEPES pH 8, 250 mM NaCl, 0.5 mM TCEP, 5% glycerol). Proteins were loaded on a HiTrap Heparin HP column (Cytiva), washed using the heparin-loading buffer, and eluted gradually from 250 mM NaCl to 1 M NaCl, over 15 CV, using 50 mM HEPES pH 8, 1 M NaCl, 2 mM TCEP, 5% glycerol. The nucleic acid-free TiLV-NP fractions (ratio A_260/280_ = 0.56) were dialyzed overnight at 4 °C in a final buffer (50 mM HEPES pH 8, 300 mM NaCl, 0.5 mM TCEP, 5% glycerol), concentrated using Amicon Ultra (10 kDa cut-off), flash-frozen in liquid nitrogen, and stored at −80 °C for further use.

### Cloning, expression and purification of ANP32A

*Oreochromis niloticus* ANP32A (UniProt A0A669BFJ9) - referred to as tilapia ANP32A (tiANP32A) - gene (TWIST) was inserted in a pETM11 plasmid using NcoI and XhoI restriction enzymes (NEB) and followed by ligation. The N-terminal His-tagged tiANP32A construct was expressed in BL21(DE3) *E. coli* cells. Expression was induced with 1 mM IPTG, for 4 h at 37 °C. Cells were harvested by centrifugation (1000 g, 20 min at 4 °C), disrupted by sonication for 5 min (5 s ON, 15 s OFF, 50% amplitude) on ice in lysis buffer (50 mM HEPES pH 8, 150 mM NaCl, 5 mM β-mercaptoethanol (BME) with cOmplete EDTA-free Protease Inhibitor Cocktail (Roche). After lysate centrifugation at 48,000 g for 45 min at 4 °C, the soluble fraction was loaded on a HisTrap HP 5 mL column (Cytiva). Bound proteins were subjected to a wash step using the lysis buffer supplemented by 50 mM imidazole. Remaining bound protein was eluted using the lysis buffer supplemented by 500 mM imidazole. Fractions containing tiANP32A were dialysed overnight in the lysis buffer (50 mM HEPES pH 8, 150 mM NaCl, 5 mM BME) together with N-terminal his-tagged TEV protease (ratio 1:5 w/w). Tag-cleaved tiANP32A protein was subjected to a Ni-sepharose affinity column to remove the TEV protease, further concentrated with Amicon Ultra-15 (3 kDa cutoff) and subjected to a size-exclusion chromatography (SEC) using a Superdex 200 Increase 10/300 GL column (Cytiva) in a final buffer containing 50 mM HEPES pH 8, 150 mM NaCl, 2 mM TCEP. Fractions containing exclusively tiANP32A were concentrated with Amicon Ultra-15 (3 kDa cutoff), flash-frozen and stored at −80 °C for later use.

Tilapia ANP32A mutants “F46A+F121A”, “F46A+F121A+E21K”, “F46A+F121A+D25K”, “F46A+F121A+I50W”, “F46A+F121A+E70K”, “F46A+F121A+E73K” were cloned using a combination of PCRs and Gibson assembly. All plasmid sequences were confirmed by Sanger sequencing. The expression and purification steps follow the same as for tiANP32A wild-type. Human ANP32A (N-terminal deca-His-tag) was expressed and purified as previously described^31^. After TEV cleavage of the His-tag, both hu and tiANP32A constructs retain an additional “GA” at the N-terminus. Truncated huANP32A constructs (1-199 and 144-249) were generated, expressed and purified as previously described^31^. The huANP32A 1-149 construct was a gift from Cynthia Wolberger (Addgene plasmid #6724151^32^) and was expressed and purified as previously described^31^.

### Analytical size-exclusion chromatography

SEC experiments for TiLV-NP and huANP32A/tiANP32A were performed on a Superdex 200 Increase

3.2/300 (Cytiva) at 4 °C, in a final buffer containing 50 mM HEPES pH 8, 150 mM NaCl, 2 mM TCEP. Depending on the experiment, 25 µM TiLV-NP were mixed with 25 µM huANP32A, 25 µM huANP32A 1-149, 25 µM huANP32A 144-249, or 25 µM tiANP32A. Resulting mixtures were incubated 1 h on ice and centrifuged 5 min at 11,000 g prior to injection onto the column. SEC fractions of interest were loaded on 4-20% Tris-glycine gel (ThermoFisher) and stained with Coomassie Blue.

### Mass photometry analysis

Mass photometry measurements were performed on a OneMP mass photometer (Refeyn). Coverslips (No. 1.5H, 24 × 50 mm, VWR) were washed with water and isopropanol before being used as a support for silicone gaskets (CultureWellTM 423 Reusable Gaskets, Grace Bio-labs). Contrast/mass calibration was realised using native marker (Native Marker unstained protein 426 standard, LC0725, Life Technologies) with a medium field of view and monitored during 60 s using the AcquireMP software (Refeyn).

For sample preparation, 0.5 µM TiLV-Pol were mixed with either 1 µM vRNA loop 40-mer, 2.5 µM huANP32A, 2.5 µM huANP32A 1-149, 2.5 µM huANP32A 1-199, 2.5 µM huANP32A 144-249, 2.5 µM tiANP32A, 2.5 µM tiANP32A “F46A+F121A”, 2.5 µM tiANP32A “F46A+F121A+E21K”, 2.5 µM tiANP32A “F46A+F121A+D25K”, 2.5 µM tiANP32A “F46A+F121A+I50W”, 2.5 µM tiANP32A “F46A+F121A+E70K”, 2.5 µM tiANP32A “F46A+F121A+E73K”.

For each condition, 18 µL of buffer (50 mM HEPES pH 8, 150 mM NaCl, 2 mM TCEP) were used to find the focus. 2 µL of sample were added to reach a final TiLV-Pol concentration of 50 nM and 250 nM ANP32A. Movies of 60 s were recorded, processed and mass estimation was determined automatically or manually using the DiscoverMP software (Refeyn).

### *In vitro* complex preparation for cryo-EM

#### TiLV polymerase in complex with tiANP32A (sample 1)

To trap apo TiLV polymerase bound to tiANP32A, 2 µM of TiLV polymerase were mixed with 10 µM of tiANP32A in the cryo-EM buffer and incubated for 1 h at 4 °C. Before proceeding to grid freezing, the sample was centrifuged for 5 min, 11,000 g and kept at 4 °C.

#### TiLV polymerase in complex with huANP32A (sample 2)

To trap apo TiLV polymerase bound to huANP32A, 2 µM of TiLV polymerase were mixed with 10 µM of huANP32A in the cryo-EM buffer and incubated for 1 h at 4 °C. Before proceeding to grid freezing, the sample was centrifuged for 5 min, 11,000 g and kept at 4 °C.

#### vRNA bound TiLV polymerase in complex with tiANP32A (sample 3)

To trap vRNA bound TiLV polymerase in complex with tiANP32A, 1.6 µM of TiLV polymerase were mixed with 1.6 µM of the TiLV 40-mer vRNA loop (5′-pGCA AAU CUU UCU CAC GUC CUG ACU UGU GAG UAA AAU UUG G −3′), 100 µM CTP and 8 µM tiANP32A in the cryo-EM buffer and incubated for 1 h at 4 °C. Before proceeding to grid freezing, the sample was centrifuged for 5 min, 11,000 g and kept at 4 °C.

### Cryo-EM grid preparation and data collection

For grid preparation, 1.5 µL of sample was applied on each sides of plasma cleaned (Fischione 1070 Plasma Cleaner: 1 min 30, 90% oxygen, 10% argon) grids (UltrAufoil 1.2/1.3, Au 300). Excess solution was blotted for 3 sec, blot force 0, 100% humidity at 4 °C with a Vitrobot Mark IV (ThermoFisher) before plunge freezing in liquid ethane.

Automated data collections were performed on a TEM Titan Krios G3 (Thermo Fisher) operated at 300 kV equipped with a K3 (Gatan) direct electron detector camera and a BioQuantum energy filter, using EPU. Coma and astigmatism correction were performed on a carbon grid. Micrographs were recorded in counting mode at a ×105,000 magnification giving a pixel size of 0.84 Å with defocus ranging from −0.8 to −2.0 µm. Gain-normalized movies of 40 frames were collected with a total exposure of ∼40 e-/Å^2^.

### Image processing

For each collected dataset, movie drift correction was performed using Relion′s Motioncor implementation, with 7 × 5 patch for K3 movies, using all movie frames^33^. All subsequent image-processing steps were performed in cryoSPARC v3.3 to v4.5^34^. CTF parameters were determined using “Patch CTF estimation”, and realigned micrographs were then inspected to manually discard low-quality images. The overall strategy to obtain all the various structures of TiLV-Pol in complex with ANP32A presented in this study is similar, with only minor variations.

Initially, particles were automatically picked using a circular blob with a diameter ranging from 90 to 140 Å. These particles were extracted using a box size of 380 × 380 pixels^2^, Fourier-cropped to 200 × 200 pixels^2^, and subjected to successive 2D classifications to remove particles with poor structural features and to roughly separate monomers from dimers. The resulting 2D classes were used for an “ab-initio reconstruction” job, after which the particles were re-extracted, Fourier-uncropped, and the best initial models used to run “non-uniform refinements”.

To separate the different conformation of monomeric TiLV-Pol, a combination of “heterogeneous refinement” and “3D classification” without alignments was applied. For each relevant TiLV-Pol conformation, a final “non-uniform refinement” was performed.

For TiLV-Pol asymmetric dimers, the corresponding particles were used for model training and picking using Topaz^35^. The newly picked particles were extracted and subjected to 2D classification. All asymmetric dimers were merged, duplicates removed, and then subjected to a “non-uniform refinement” job. To improve the density of the respective TiLV-Pol replicase and ANP32-encapsidase moieties, particle subtraction around ′TiLV-Pol(R) minus 627(R)′ and ′TiLV-Pol(E)-ANP32A-627(R)′ were performed. These subtracted particles were subjected to “local refinements”. Based on these consensus maps, further “3D classifications” without alignments were used to separate the different conformations of TiLV-Pol replicase and encapsidase. A final “local refinement” job was performed for each conformation of interest.

Post-processing was performed in cryoSPARC using an automatically or manually determined B-factor. For each final map, reported global resolution is based on the FSC 0.143 cut-off criteria. Local resolution variations were estimated in cryoSPARC.

For more detailed image processing information regarding each data collection and deposited maps, please refer to Fig. S2-S8.

### Model building and refinement

Atomic models of the TiLV replication complexes and TiLV encapsidase bound to tiANP32A and huANP32A were constructed by iterative rounds of manual model building within COOT^36^. For model building of the replicase-moiety of the TiLV replication complex, the previously determined partial replicase structure (PDB 8PT6)^7^ was used as starting point. The PB2/627 domain was extracted from the transcriptase structure (PDB 8PSZ)^5^ and rigidly fitted into the density for subsequent adjustments. For the TiLV-Pol encapsidase, similar procedure has been applied to fit the PB2/627-NLS and PA/ENDO domains. For huANP32A and tiANP32A, the crystal structure of the LRR domain (PDB 2JE1)^37^ and the Alphafold3^38^ structure prediction were used as a starting point, respectively.

Models were refined using Phenix real-space refinement^39^ with Ramachandran restraints. Atomic model validation was performed using Molprobity^40^ as implemented in Phenix^41^. Model resolution according to cryo-EM map was estimated at the 0.5 FSC cutoff. Buried solvent accessible surfaces were calculated using PISA^42^ at the PDBe. Electrostatic potential was calculated using PDB2PQR^43^ and APBS^44^. Fig.s were generated using ChimeraX^45^.

## Acknowledgments

We acknowledge the European Synchrotron Radiation Facility (ESRF) and the Partnership for Structural Biology (PSB) for access to the Titan Krios CM01, and especially Eaazhisai Kandiah for setting up the multiple high-quality cryo-EM data collections. We thank Romain Linares and Iskander Khusainov for access to the EMBL Grenoble Glacios. Aymeric Peuch for support using the joint EMBL-IBS computer cluster; Caroline Mas for assistance and access to the biophysical platform. This work used the platforms of the Grenoble Instruct-ERIC center (ISBG; UAR 3518 CNRS-CEA-UGA-EMBL) within the Grenoble Partnership for Structural Biology (PSB), supported by FRISBI (ANR-10-INBS-0005-02) and GRAL, financed within the University Grenoble Alpes graduate school (Ecoles Universitaires de Recherche) CBH-EUR-GS (ANR-17-EURE-0003).

## Author contributions

SC and BA conceived the project. BA and MP performed cloning. BA performed protein expression, and purification. BA performed all *in vitro* biochemical, biophysical and cryo-EM analyses. SC and BA did model building and refinement. BA and SC prepared the manuscript.

## Data, code, and materials availability

The coordinates and EM maps generated in this study have been deposited in the Protein Data Bank and the Electron Microscopy Data Bank (see Table 1) under accession codes:

PDB 33FF / EMD-59408: Monomeric apo TiLV polymerase in encapsidase conformation bound to tilapia ANP32A (open core);

PDB 33FG / EMD-59409: TiLV replication complex bound to tilapia ANP32A: Local refinement around the replicase minus 627(R);

PDB 33FH / EMD-59410: TiLV replication complex bound to tilapia ANP32A: Local refinement around the encapsidase-tiANP32A plus 627(R);

PDB 33FQ / EMD-59419: TiLV replication complex bound to tilapia ANP32A (consensus refinement);

PDB 33FI / EMD-59411: Monomeric apo TiLV polymerase in encapsidase conformation bound to human ANP32A (open core);

PDB 33FJ / EMD-59412: TiLV replication complex bound to human ANP32A: Local refinement around the replicase minus 627(R);

PDB 33FK / EMD-59413: TiLV replication complex bound to human ANP32A: Local refinement around the encapsidase-huANP32A plus 627(R);

PDB 33FR / EMD-59420: TiLV replication complex bound to human ANP32A (consensus refinement);

PDB 33FL / EMD-59414: vRNA-bound TiLV replication complex bound to tilapia ANP32A: Local refinement around the encapsidase-tiANP32A plus 627(R) in initiation state (mode A);

PDB 33FM / EMD-59415: vRNA-bound TiLV replication complex bound to tilapia ANP32A: Local refinement around the encapsidase-tiANP32A plus 627(R) with 5′ hook bound;

PDB 33FN / EMD-59416: vRNA-bound TiLV replication complex bound to tilapia ANP32A: Local refinement around the replicase minus 627(R) in initiation state (mode A);

PDB 33FS / EMD-59421: vRNA-bound TiLV replication complex bound to tilapia ANP32A: Consensus refinement with the replicase in initiation mode A;

PDB 33FO / EMD-59417: vRNA-bound TiLV replication complex bound to tilapia ANP32A: Local refinement around the replicase minus 627(R) in mode B with closed core;

PDB 33FT / EMD-59422: vRNA-bound TiLV replication complex bound to tilapia ANP32A: Consensus refinement with the replicase in mode B with closed core;

PDB 33FP / EMD-59418: vRNA-bound TiLV replication complex bound to tilapia ANP32A: Local refinement around the replicase minus 627(R) in mode B with open core;

PDB 33FU / EMD-59423: vRNA-bound TiLV replication complex bound to tilapia ANP32A: Consensus refinement with the replicase in mode B with open core.

## SUPPORTING INFORMATION CAPTIONS

**Fig. S1. Multiple sequence alignment of fish, human, and chicken ANP32 variants.**

The alignment includes ANP32 sequences from *Oreochromis niloticus* (tilapia; UniProt A0A669BFJ9), *Danio rerio* (zebrafish; UniProt Q7ZUP0), *Salmo salar* (Atlantic salmon; UniProt A0A1S3QKC6), *Homo sapiens* (human) ANP32A (UniProt P39687), ANP32B (UniProt Q92688), and ANP32E (UniProt Q9BTT0), as well as *Gallus gallus* (chicken) ANP32A (UniProt XP_413932.3). The ANP32 leucine-rich repeat (LRR) and low-complexity acidic region (LCAR) domains are indicated above the sequences in dark and light purple, respectively. Residues of tiANP32A that interact with the TiLV-Pol encapsidase are marked below the alignment with coloured circles corresponding to contacts with the TiLV-Pol PB2/627(E) (light pink), PB2/NLS(E) (brown), PB2/627(R) (pink) and PA-C(E) (light green) domains, respectively. The 33-amino acid insertion specific to avian ANP32A is highlighted by a grey rectangle.

**Fig. S2. Cryo-EM image processing strategy applied to obtain TiLV polymerase structures bound to tiANP32A.**

Schematics of the image processing strategy used with the data collected on a Titan Krios equipped with a Gatan K3 direct electron detector mounted on a Gatan Bioquantum energy filter. Representative micrograph, 2D class averages and 3D class averages are displayed. Views of each sharpened EM maps are shown. Fourier shell correlation curves (FSC) are displayed.

**Fig. S3. Local resolution, cFSC curves, and particle orientation distributions for TiLV polymerase structures bound to tiANP32A.**

Sharpened cryo-EM maps are coloured according to estimated local resolution (Å), alongside their respective conical Fourier shell correlation (cFSC) curves and particle orientation distribution plots. (A) Monomeric apo TiLV polymerase in encapsidase conformation bound to tilapia ANP32A (open core). (B) TiLV replication complex bound to tilapia ANP32A (consensus refinement). (C) TiLV replication complex bound to tilapia ANP32A: Local refinement around the replicase minus 627(R). (D) TiLV replication complex bound to tilapia ANP32A: Local refinement around the encapsidase-tiANP32A plus 627(R).

**Fig. S4. Cryo-EM image processing strategy applied to obtain TiLV polymerase structures bound to huANP32A.**

Schematics of the image processing strategy used with the data collected on a Titan Krios equipped with a Gatan K3 direct electron detector mounted on a Gatan Bioquantum energy filter. Representative micrograph, 2D class averages and 3D class averages are displayed. Views of each sharpened EM maps are shown. Fourier shell correlation curves (FSC) are displayed.

**Fig. S5. Local resolution, cFSC curves, and particle orientation distributions for TiLV polymerase structures bound to huANP32A.**

Sharpened cryo-EM maps are coloured according to estimated local resolution (Å), alongside their respective conical Fourier shell correlation (cFSC) curves and particle orientation distribution plots. (**A**) Monomeric apo TiLV polymerase in encapsidase conformation bound to human ANP32A (open core). (**B**) TiLV replication complex bound to human ANP32A (consensus refinement). (**C**) TiLV replication complex bound to human ANP32A: Local refinement around the replicase minus 627(R). (**D**) TiLV replication complex bound to human ANP32A: Local refinement around the encapsidase-huANP32A plus 627(R).

**Fig. S6. Cryo-EM image processing strategy applied to obtain vRNA-bound TiLV polymerase structures in complex with tiANP32A (focus on the replicase).**

Schematics of the image processing strategy used with the data collected on a Titan Krios equipped with a Gatan K3 direct electron detector mounted on a Gatan Bioquantum energy filter. Representative micrograph, 2D class averages and 3D class averages are displayed. Views of each sharpened EM maps are shown. Fourier shell correlation curves (FSC) are displayed.

**Fig. S7. Cryo-EM image processing strategy applied to obtain vRNA-bound TiLV polymerase structures in complex with tiANP32A (focus on the encapsidase).**

Schematics of the image processing strategy used with the data collected on a Titan Krios equipped with a Gatan K3 direct electron detector mounted on a Gatan Bioquantum energy filter. Representative micrograph, 2D class averages and 3D class averages are displayed. Views of each sharpened EM maps are shown. Fourier shell correlation curves (FSC) are displayed.

**Fig. S8. Local resolution, cFSC curves, and particle orientation distributions for vRNA-bound TiLV polymerase structures in complex with tiANP32A.**

Sharpened cryo-EM maps are coloured according to estimated local resolution (Å), alongside their respective conical Fourier shell correlation (cFSC) curves and particle orientation distribution plots. (**A**) vRNA-bound TiLV replication complex bound to tilapia ANP32A: Local refinement around the encapsidase-tiANP32A plus 627(R) in initiation state (mode A). (**B**) vRNA-bound TiLV replication complex bound to tilapia ANP32A: Local refinement around the encapsidase-tiANP32A plus 627(R) with 5′ hook bound. (**C**) vRNA-bound TiLV replication complex bound to tilapia ANP32A: Local refinement around the replicase minus 627(R) in initiation state (mode A). (**D**) vRNA-bound TiLV replication complex bound to tilapia ANP32A: Local refinement around the replicase minus 627(R) in mode B with closed core. (**E**) vRNA-bound TiLV replication complex bound to tilapia ANP32A: Local refinement around the replicase minus 627(R) in mode B with open core. (**F**) vRNA-bound TiLV replication complex bound to tilapia ANP32A: Local refinement around the replicase minus 627(R) in mode B with distal duplex.

**Fig. S9. Structural comparison of the monomeric apo TiLV polymerase in encapsidase conformation.**

(**A**) Cut-away view of the sharpened cryo-EM map for the monomeric apo TiLV polymerase in encapsidase conformation (TiLV-Pol(E)) bound to tilapia ANP32A, focused on the main interface between tiANP32A, PB2/627-NLS(E), PB1(E), and PB2-N(E). Side-chain densities are clearly visible. tiANP32A is coloured purple, PB2/NLS is brown, PB1(E) is light blue, and PB2-N is dark grey. Water molecules are shown as red spheres. (**B**) Cut-away view of the sharpened cryo-EM map for the monomeric apo TiLV polymerase in encapsidase conformation (TiLV-Pol(E)) bound to human ANP32A. Domains and map densities are coloured as in panel **A**. (**C**) Superposition of the PB2/627-NLS domains from the huANP32A-bound and tiANP32A-bound TiLV-Pol(E) structures reveals a slight structural shift of ∼6 Å for the ANP32A LRR domain. The RMSD between the respective PB2/627-NLS domains is ∼0.3 Å, while the RMSD between the interacting residues of ANP32A is ∼2.4 Å. (**D**) View of all TiLV Pol(E)-ANP32A interacting residues reveals a highly conserved mode of interaction between the human and tilapia variants.

**Fig. S10. Interfaces of the PB2/627-NLS(E) domain with PA-C(E) and PB1(E) block the access to the secondary site.**

(**A**) Interface between PB2/627-NLS(E) and PA-C(E). Top: Exploded view of the tiANP32A-PB2/627-NLS(E) subcomplex and the TiLV-Pol(E). Structures are shown as surfaces. The PB2/627(E) footprint on PA-C(E) is highlighted in light pink, and conversely, the PA-C(E) footprint on PB2/627(E) is coloured light green. Domains are coloured as follows: PA is light green, PB1 is light blue, PB2-N is dark grey, PB2/627 is light pink, PB2/NLS is brown, and tiANP32A is purple. Bottom: Detailed molecular interactions between PA-C(E) and PB2/627(E). Interacting residues are shown as sticks. Hydrogen bonds and salt bridges are indicated by dotted lines. (**B**) Interface between PB2/627-NLS(E) and PB1(E). Top: Exploded view of the tiANP32A-PB2/627-NLS(E) subcomplex and the TiLV-Pol(E). Structures are shown as surfaces. The PB2/627-NLS(E) footprint on PB1(E) is highlighted, and conversely, the corresponding PB1(E) footprint is displayed on PB2/627-NLS(E). Domains are coloured as in panel **A**. Bottom: Detailed molecular interactions between PB1(E), PB2/627(E) and PB2/NLS(E). Interacting residues are shown as sticks. Hydrogen bonds and salt bridges are indicated by dotted lines. (**C**) Close-up view of TiLV-Pol(T) in pre-termination state (PDB 8QZ8), with the 3′ vRNA end occupying the secondary binding site. The structure is shown as a surface with the underlying ribbon representation visible. PA-C(T) is dark green, PB1(T) is cyan, PB2-N(T) is bordeaux, and the 3′ vRNA is gold. (**D**) Superposition of the 3′ vRNA end onto the monomeric apo TiLV-Pol(E) bound to tilapia ANP32A reveals that the secondary binding site is sterically occluded. Structures were aligned using PA-C as a reference. Residues clashing with the 3′ vRNA end (distance < 0.6 Å) are coloured red and labelled. Domains are coloured as in panels **A** and **B**.

**Fig. S11. Structural comparison between the three polymerase functionnal conformations observed within the *Articulavirales* order.**

(**A-C**) Structural comparison of the PB2/627-NLS domains across the (**A**) TiLV-Pol transcriptase (T), (**B**) TiLV-Pol replicase (R), and (**C**) TiLV-Pol encapsidase (E) conformations. Domains are coloured as follows: PB2/NLS is beige, PB2/627 is pink, PA/ENDO is green, PB2/Mid-link is dark pink, and tiANP32A is purple. The PB2/627-NLS linker (residues 375-382) is highlighted with a black solid line. (**D-F**) All structures are displayed in cartoon representation. The polymerase core is coloured dark grey, while domains coloured as follow: PA/ENDO (or ENDO-like) in green, PB2/CBD (or CBD-like) in orange, PB2/Mid-link in purple, PB2/627 in pink, PB2/NLS in beige, and ANP32A in purple. Where present, the 5′ and 3′ vRNA promoter ends are shown transparent in plum and gold, respectively. The PA-C domain is indicated for the encapsidase conformations. (**D**) Monomeric TiLV-Pol in transcriptase (T) conformation (PDB 8PSN) and FluPol (A/H17N10) in transcriptase conformation (PDB 4WSB). (**E**) Left: Monomeric TiLV-Pol in partial replicase (R) conformation (PDB 8PT6). The PB2/627 domain remains flexible and is unresolved when not part of the fully assembled replication complex. Right: TiLV-Pol in complete replicase conformation, extracted from the TiLV replication complex. Bottom: FluPol (A/H7N9) in complete replicase conformation, extracted from the FluA replication complex (PDB 8RMR). (**F**) Top: Monomeric, tiANP32A-bound apo-TiLV-Pol in encapsidase (E) conformation. The PB2/Mid-link and PB2/CBD domains are not visible due to flexibility. Bottom left: FluPol (A/H7N9) in complete replicase conformation, extracted from the FluA replication complex (PDB 8RMR). Bottom right: Monomeric, huANP32B-bound FluPol (A/H5N1) in partial encapsidase conformation (PDB 8R1L).

**Fig. S12. Specific tiANP32A mutations disrupt TiLV replication complex formation in vitro.**

(**A**) View of the tiANP32A binding site on the tripartite 627(E)-NLS(E)-627(R) complex. PB2/627(E), PB2/NLS(E), and PB2/627(R) are shown as surfaces and coloured light pink, brown, and pink, respectively. tiANP32A is shown in cartoon representation, with the mutated residues highlighted by black lines and displayed as sticks. (**B-G**) Mass photometry analysis of TiLV-Pol mixed with (**B**) the tiANP32A F46A/F121A double mutant (DM), (**C**) tiANP32A DM+E21K, (**D**) tiANP32A DM+D25K, (**E**) tiANP32A DM+I50W, (**F**) tiANP32A DM+E70K, and (**G**) tiANP32A DM+E73K. Plots display normalized counts versus mass (kDa). Solid black lines indicate the main peaks. The estimated mass and relative percentage for each population. Asterisks (*) indicate putative dimers that could not be precisely measured.

**Fig. S13. Structural comparison of replication complexes across the *Articulavirales* order.**

(**A-E**) Comparative views of replication complexes from the *Amnoonviridae* (**A**) and *Orthomyxoviridae* (**B-E**) families. For each panel, three structural representations are provided: Left: Close-up view of the PA/ENDO(R) and PB2/NLS(R) interaction. Structures are aligned on the PA/ENDO domain. Amino acid boundaries for the respective domains are indicated. Middle: Overall architecture of the replication complex. Structures are aligned on the PB1(R). Right: Close-up view of the host-virus tripartite 627(R)- NLS(E)-627(E) interface induced by the ANP32 LRR domain. (**A**) TiLV replication complex bound to tiANP32A. (**B**) Influenza A (FluA) replication complex bound to huANP32A (PDB 8RMR). (**C**) Influenza B (FluB) replication complex bound to huANP32A (PDB 8RNC). (**D**) Influenza C (FluC) replication complex bound to chANP32A (PDB 6XZR). The FluC replication complex was observed with/without bound ANP32. (**E**) Thogoto virus (THOV) replication complex-like structure (PDB 8Z9R). Due to the absence of a host ANP32 factor, the PB2/627-NLS(E) remains flexible and unresolved. A putative binding site for a tick ANP32 variant is indicated by a purple arrow.

**Fig. S14. TiLV-NP interacts with the ANP32A LCAR in vitro.**

(**A-E**, **G-J**) SDS-PAGE analysis of size-exclusion chromatography (SEC) fractions for: (**A**) TiLV-NP alone (the two displayed gels are duplicated for visual purpose), (**B**) tilapia ANP32A (tiANP32A) alone, (**C**) human ANP32A (huANP32A) alone, (**D**) TiLV-NP incubated with tiANP32A, (**E**) TiLV-NP with huANP32A, (**G**) the huANP32A LRR domain alone, (**H**) the huANP32A LCAR domain alone, (**I**) TiLV-NP with the huANP32A LRR domain, and (**J**) TiLV-NP with the huANP32A LCAR domain. For all gels, molecular weight (MW) markers (in kDa) and the proteins are indicated. Elution fraction numbers are indicated above each lane. (**F**) Superposition of SEC elution profiles for TiLV-NP alone (solid grey line), TiLV-NP incubated with ANP32A (solid black lines), and ANP32A alone (solid purple lines), with all main peaks annotated accordingly. Relative absorbance at 280 nm (mAU) is plotted on the y-axis, and elution volume (mL) is plotted on the x-axis (graduated every 50 µL). (**K**) Superposition of SEC elution profiles for TiLV-NP alone (solid grey line), TiLV-NP with the huANP32A LRR domain (dotted black line), TiLV-NP with the huANP32A LCAR domain (solid black line), and the isolated LCAR and LRR domains alone (solid purple lines), with all main peaks annotated accordingly. Axes are formatted identical to panel **F**.

**Table S1.** Cryo-EM data collection, refinement and validation statistics for the monomeric TiLV encapsidase and the apo TiLV replication complex, bound to tiANP32A.

| TiLV polymerase in complex with tiANP32A |  |  |  |  |
| --- | --- | --- | --- | --- |
| No. | 1 | 2 | 3 | 4 |
| Structure | Monomeric apo TiLV polymerase in encapsidase conformation bound to tilapia ANP32A (open core) | TiLV replication complex bound to tilapia ANP32A: Local refinement around the replicase minus 627(R) | TiLV replication complex bound to tilapia ANP32A: Local refinement around the encapsidase-tiANP32A plus 627(R) | TiLV replication complex bound to tilapia ANP32A (consensus refinement) |
| PDB ID | 33FF | 33FG | 33FH | 33FQ |
| EMDB ID | EMD-59408 | EMD-59409 | EMD-59410 | EMD-59419 |
| Data collection and processing |  |  |  |  |
| Microscope | ThermoFisher Krios TEM |  |  |  |
| Voltage (kV) | 300 |  |  |  |
| Camera | Gatan K3 direct electron detector mounted on a Gatan Bioquantum LS/967 energy filter |  |  |  |
| Magnification | 105,000 |  |  |  |
| Nominal defocus range (μm) | -0.8 / -2 |  |  |  |
| Electron exposure (e<sup>-</sup>/Å<sup>2</sup>) | 40 |  |  |  |
| Pixel size (Å) | 0.84 |  |  |  |
| Initial micrographs (no.) | 8,720 |  |  |  |
| Final micrographs (no.) | 7,432 |  |  |  |
| Refinement |  |  |  |  |
| Particles per class (no.) | 263,306 | 34,806 |  |  |
| Map resolution (Å), 0.143 FSC | 2.69 | 3.10 | 3.05 | 3.17 |
| Model resolution (Å), 0.5 FSC | 2.8 | 3.4 | 3.4 | 3.5 |
| Map sharpening *B* factor (Å<sup>2</sup>) | -96.6 | -75.1 | -71.3 | -75.9 |
| Map versus model cross-correlation (CCmask) | 0.8294 | 0.7666 | 0.7792 | 0.7479 |
| Model composition |  |  |  |  |
| Non-hydrogen atoms | 10357 | 10117 | 11642 | 21217 |
| Protein residues | 1311 | 1268 | 1481 | 2701 |
| Nucleotides | - | 8 | - | - |
| Water | 60 | 1 | - | - |
| Ligands | 1xZn | 3xZn | 1xZn | 4xZn |
| *B* factors (Å<sup>2</sup>) |  |  |  |  |
| Protein | 55.47 | 107.03 | 101.90 | 140.52 |
| RNA | - | 143.01 | - | - |
| Water | 27.55 | 241.98 | - | - |
| Ligands | 57.02 | 69/99 | 107.89 | 259.67 |
| RMS deviations |  |  |  |  |
| Bond lengths (Å) | 0.002 | 0.002 | 0.003 | 0.002 |
| Bond angles (°) | 0.454 | 0.435 | 0.422 | 0.432 |
| Validation |  |  |  |  |
| MolProbity score | 1.36 | 2.16 | 1.75 | 1.74 |
| All-atom clash score | 1.45 | 5.66 | 4.24 | 4.73 |
| Poor rotamers (%) | 2.97 | 5.59 | 2.63 | 1.26 |
| Ramachandran plot |  |  |  |  |
| Favored (%) | 97.37 | 95.87 | 96.50 | 96.56 |
| Allowed (%) | 2.63 | 3.82 | 3.50 | 3.33 |
| Outliers (%) | 0.0 | 0.32 | 0.0 | 0.11 |

**Table S2.** Cryo-EM data collection, refinement and validation statistics for the monomeric TiLV encapsidase and the apo TiLV replication complex, bound to huANP32A.

| TiLV polymerase in complex with huANP32A |  |  |  |  |
| --- | --- | --- | --- | --- |
| No. | 5 | 6 | 7 | 8 |
| Structure | Monomeric apo TiLV polymerase in encapsidase conformation bound to human ANP32A (open core) | TiLV replication complex bound to human ANP32A: Local refinement around the replicase minus 627(R) | TiLV replication complex bound to human ANP32A: Local refinement around the encapsidase-huANP32A plus 627(R) | TiLV replication complex bound to human ANP32A (consensus refinement) |
| PDB ID | 33FI | 33FJ | 33FK | 33FR |
| EMDB ID | EMD-59411 | EMD-59412 | EMD-59413 | EMD-59420 |
| Data collection and processing |  |  |  |  |
| Microscope | ThermoFisher Krios TEM |  |  |  |
| Voltage (kV) | 300 |  |  |  |
| Camera | Gatan K3 direct electron detector mounted on a Gatan Bioquantum LS/967 energy filter |  |  |  |
| Magnification | 105,000 |  |  |  |
| Nominal defocus range (μm) | -0.8 / -2 |  |  |  |
| Electron exposure (e-/Å²) | 40 |  |  |  |
| Pixel size (Å) | 0.84 |  |  |  |
| Initial micrographs (no.) | 15,620 |  |  |  |
| Final micrographs (no.) | 10,817 |  |  |  |
| Refinement |  |  |  |  |
| Particles per class (no.) | 116,424 | 69,318 |  |  |
| Map resolution (Å), 0.143 FSC | 2.99 | 3.18 | 3.00 | 3.15 |
| Model resolution (Å), 0.5 FSC | 3.1 | 3.4 | 3.3 | 3.4 |
| Map sharpening *B* factor (Å²) | -110.4 | -99.1 | -93.8 | -94.1 |
| Map versus model cross-correlation (CCmask) | 0.8452 | 0.7996 | 0.8051 | 0.7869 |
| Model composition |  |  |  |  |
| Non-hydrogen atoms | 10677 | 10142 | 11540 | 21182 |
| Protein residues | 1354 | 1270 | 1471 | 2698 |
| Nucleotides | - | 8 | - | - |
| Water | 9 | - | - | - |
| Ligands | 1xZn | 3xZn, 1xMg | 1xZn | 4xZn |
| *B* factors (Å²) |  |  |  |  |
| Protein | 63.96 | 92.08 | 96.45 | 133.12 |
| RNA | - | 136.26 | - | - |
| Water | 20.52 | - | - | - |
| Ligands | 84.67 | 182.26 | 101.52 | 244.02 |
| RMS deviations |  |  |  |  |
| Bond lengths (Å) | 0.002 | 0.002 | 0.002 | 0.003 |
| Bond angles (°) | 0.428 | 0.429 | 0.438 | 0.440 |
| Validation |  |  |  |  |
| MolProbity score | 1.75 | 1.89 | 1.81 | 1.94 |
| All-atom clash score | 3.41 | 3.57 | 44.5 | 6.06 |
| Poor rotamers (%) | 3.62 | 2.93 | 2.22 | 3.12 |
| Ramachandran plot |  |  |  |  |
| Favored (%) | 96.86 | 96.19 | 97.46 | 96.44 |
| Allowed (%) | 3.06 | 3.57 | 2.54 | 3.41 |
| Outliers (%) | 0.07 | 0.24 | 0.0 | 0.15 |

**Table S3.** Cryo-EM data collection, refinement and validation statistics for the vRNA bound TiLV replication complex, bound to tiANP32A.

| vRNA bound TiLV replication complex (with tiANP32A) |  |  |  |  |  |  |  |  |
| --- | --- | --- | --- | --- | --- | --- | --- | --- |
| No. | 9 | 10 | 11 | 12 | 13 | 14 | 15 | 16 |
| Structure | vRNA-bound TiLV replication complex bound to tilapia ANP32A: Local refinement around the encapsidase-tiANP32A plus 627(R) in initiation state (mode A) | vRNA-bound TiLV replication complex bound to tilapia ANP32A: Local refinement around the encapsidase-tiANP32A plus 627(R) with 5' hook bound | vRNA-bound TiLV replication complex bound to tilapia ANP32A: Local refinement around the replicase minus 627(R) in initiation state (mode A) | vRNA-bound TiLV replication complex bound to tilapia ANP32A: Consensus refinement with the replicase in initiation mode A | vRNA-bound TiLV replication complex bound to tilapia ANP32A: Local refinement around the replicase minus 627(R) in mode B with closed core | vRNA-bound TiLV replication complex bound to tilapia ANP32A: Consensus refinement with the replicase in mode B with closed core | vRNA-bound TiLV replication complex bound to tilapia ANP32A: Local refinement around the replicase minus 627(R) in mode B with open core | vRNA-bound TiLV replication complex bound to tilapia ANP32A: Consensus refinement with the replicase in mode B with open core |
| PDB ID | 33FL | 33FM | 33FN | 33FS | 33FO | 33FT | 33FP | 33FU |
| EMDB ID | EMD-59414 | EMD-59415 | EMD-59416 | EMD-59421 | EMD-59417 | EMD-59422 | EMD-59418 | EMD-59423 |
| Data collection and processing |  |  |  |  |  |  |  |  |
| Microscope | ThermoFisher Krios TEM |  |  |  |  |  |  |  |
| Voltage (kV) | 300 |  |  |  |  |  |  |  |
| Camera | Gatan K3 direct electron detector mounted on a Gatan Bioquantum LS/967 energy filter |  |  |  |  |  |  |  |
| Magnification | 105,000 |  |  |  |  |  |  |  |
| Nominal defocus range (μm) | -0.8 / -2 |  |  |  |  |  |  |  |
| Electron exposure (e-/Å²) | 40 |  |  |  |  |  |  |  |
| Pixel size (Å) | 0.84 |  |  |  |  |  |  |  |
| Initial micrographs (no.) | 11,442 |  |  |  |  |  |  |  |
| Final micrographs (no.) | 7,922 |  |  |  |  |  |  |  |
| Refinement |  |  |  |  |  |  |  |  |
| Particles per class (no.) | 344,945 | 57,230 | 58,206 |  | 392,620 |  | 126,018 |  |
| Map resolution (Å), 0.143 FSC | 2.45 | 2.84 | 2.83 | 3.09 | 2.51 | 2.64 | 2.71 | 2.87 |
| Model resolution (Å), 0.5 FSC | 2.7 | 3.2 | 3.1 | 3.4 | 2.7 | 2.8 | 3.1 | 3.2 |
| Map sharpening *B* factor (Å²) | -77.9 | -65.5 | -72.3 | -78.1 | -83 | -92.7 | -76.4 | -80.6 |
| Map versus model cross-correlation (CCmask) | 0.8153 | 0.7634 | 0.8009 | 0.7440 | 0.8029 | 0.7754 | 0.7902 | 0.7360 |
| Model composition |  |  |  |  |  |  |  |  |
| Non-hydrogen atoms | 11688 | 11861 | 10462 | 22075 | 10153 | 21716 | 10462 | 22000 |
| Protein residues | 1384 | 1486 | 1245 | 2628 | 1251 | 2619 | 1255 | 2629 |
| Nucleotides | 31 | 8 | 29 | 60 | 14 | 45 | 28 | 59 |
| Water | 62 | 0- | 1 | 0 | 26 | 88 | 1 | - |
| Ligands | 1xZn, 1xMg, 1xCTP | 1xZn | 3xZn, 1xMg, 1xCTP | 4xZn, 2xMg, 2xCTP | 3xZn | 4xZn, 1xMg | 3xZn | 4xZn, 1xMg, 1xCTP |
| *B* factors (Å²) |  |  |  |  |  |  |  |  |
| Protein | 67.85 | 88.50 | 96.05 | 120.12 | 73.14 | 90.89 | 88.72 | 105.57 |
| RNA | 92.02 | 102.09 | 126.73 | 170.98 | 78.24 | 126.69 | 132.95 | 179.66 |
| Water | 68.20 | - | 55.19 | - | 52.86 | 69.50 | 69.01 | - |
| Ligands | 55.44 | 95.13 | 182.96 | 197.35 | 163.58 | 162.18 | 194.60 | 177.47 |
| RMS deviations |  |  |  |  |  |  |  |  |
| Bond lengths (Å) | 0.003 | 0.002 | 0.003 | 0.002 | 0.002 | 0.002 | 0.003 | 0.003 |
| Bond angles (°) | 0.484 | 0.416 | 0.411 | 0.436 | 0.429 | 0.430 | 0.439 | 0.425 |
| Validation |  |  |  |  |  |  |  |  |
| MolProbity score | 2.99 | 1.37 | 1.91 | 1.65 | 1.71 | 1.46 | 1.76 | 1.73 |
| All-atom clash score | 4.47 | 1.77 | 4.04 | 4.09 | 3.53 | 3.78 | 3.83 | 4.19 |
| Poor rotamers (%) | 3.23 | 2.00 | 4.01 | 2.32 | 2.59 | 1.50 | 2.59 | 1.97 |
| Ramachandran plot |  |  |  |  |  |  |  |  |
| Favored (%) | 97.01 | 96.74 | 96.19 | 96.93 | 96.28 | 97.07 | 95.97 | 95.67 |
| Allowed (%) | 2.99 | 3.19 | 3.81 | 3.03 | 3.64 | 2.89 | 4.03 | 4.25 |
| Outliers (%) | 0.0 | 0.07 | 0.0 | 0.04 | 0.08 | 0.04 | 0.0 | 0.08 |

