## Supplemental Figures for "Structures of the Tilapia Lake Virus replication complex show that ANP32 is a host factor for replication across the *Articulavirales* order"

### SUPPLEMENTARY INFORMATION 1

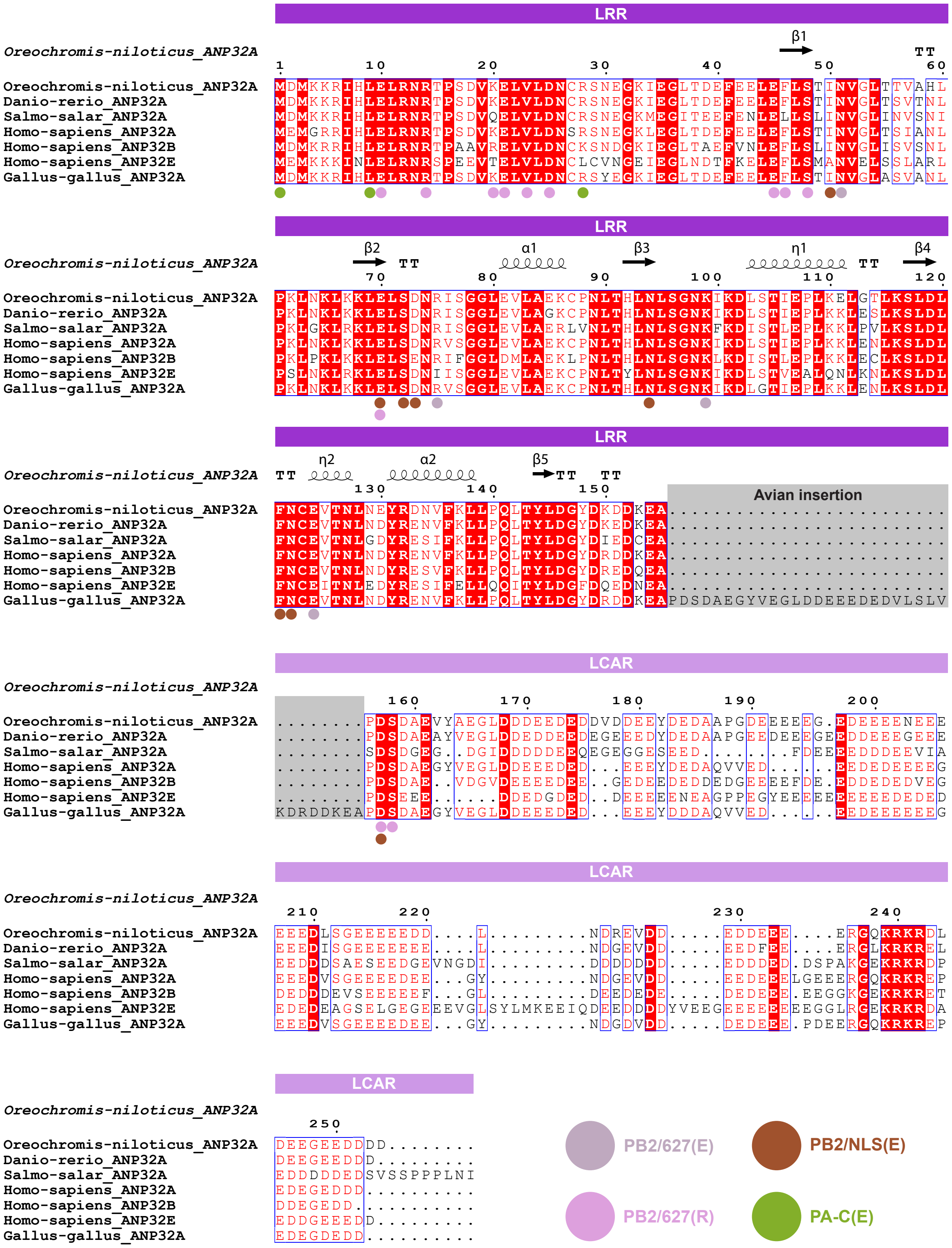

### SUPPLEMENTARY INFORMATION 2

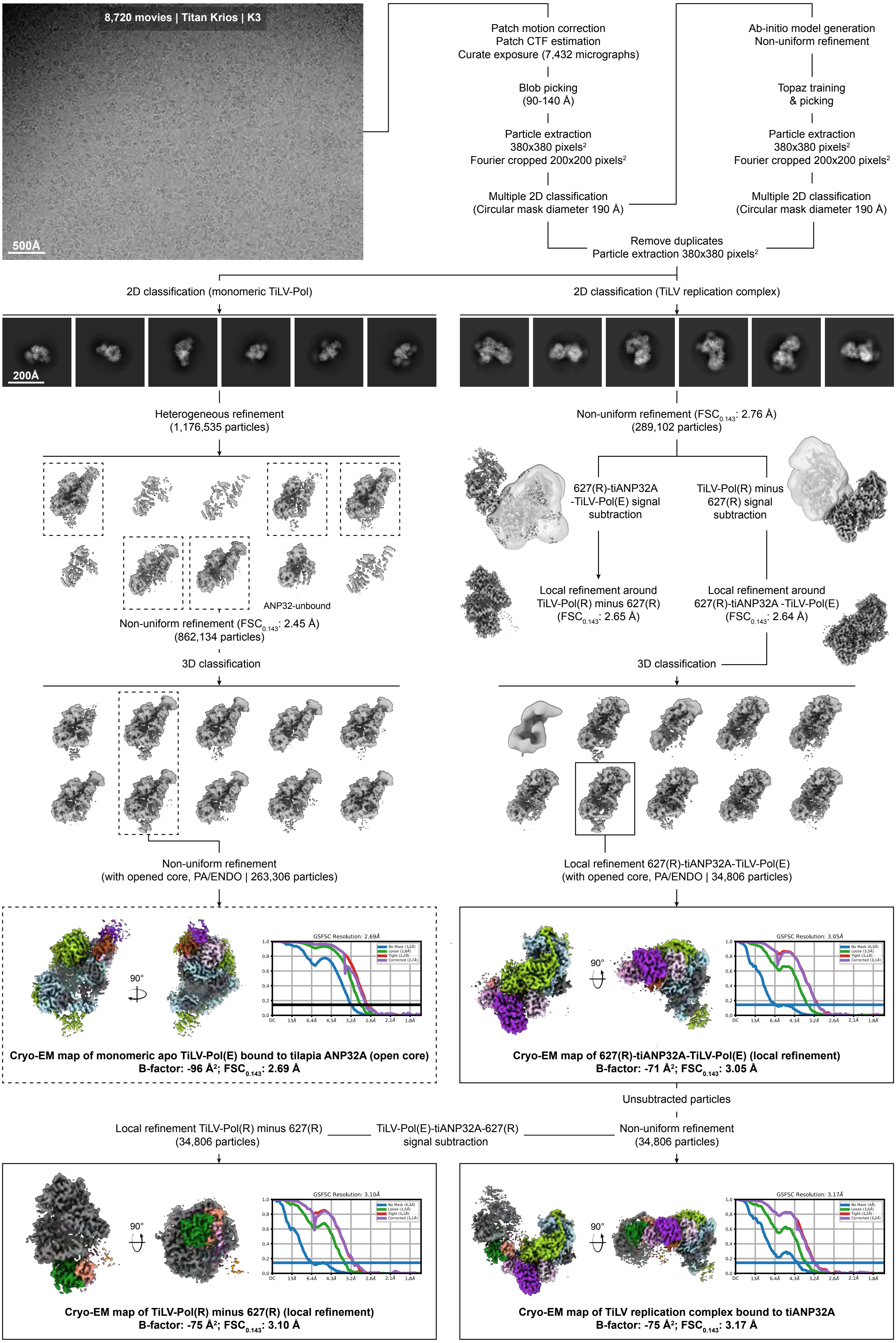

### SUPPLEMENTARY INFORMATION 3

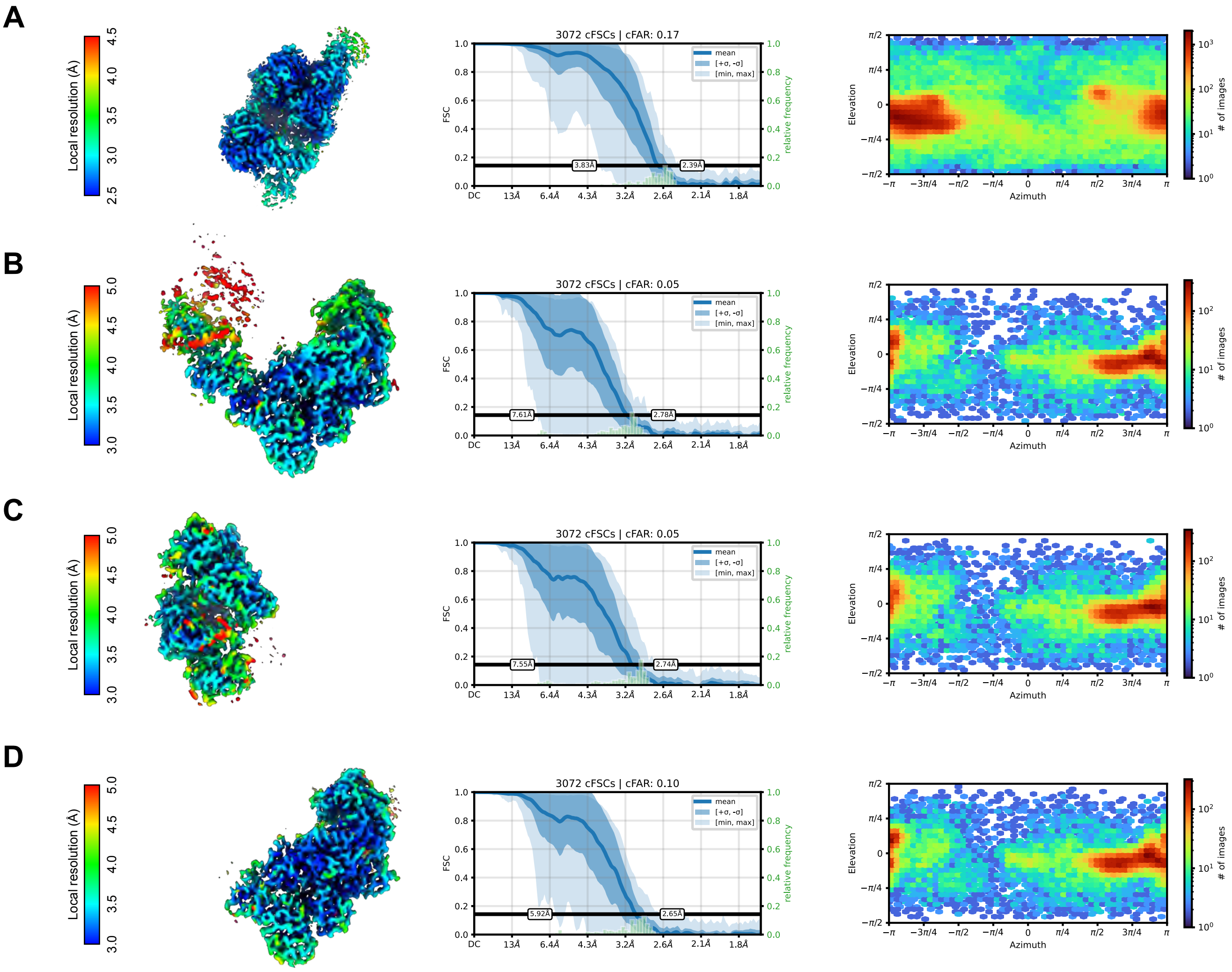

### SUPPLEMENTARY INFORMATION 4

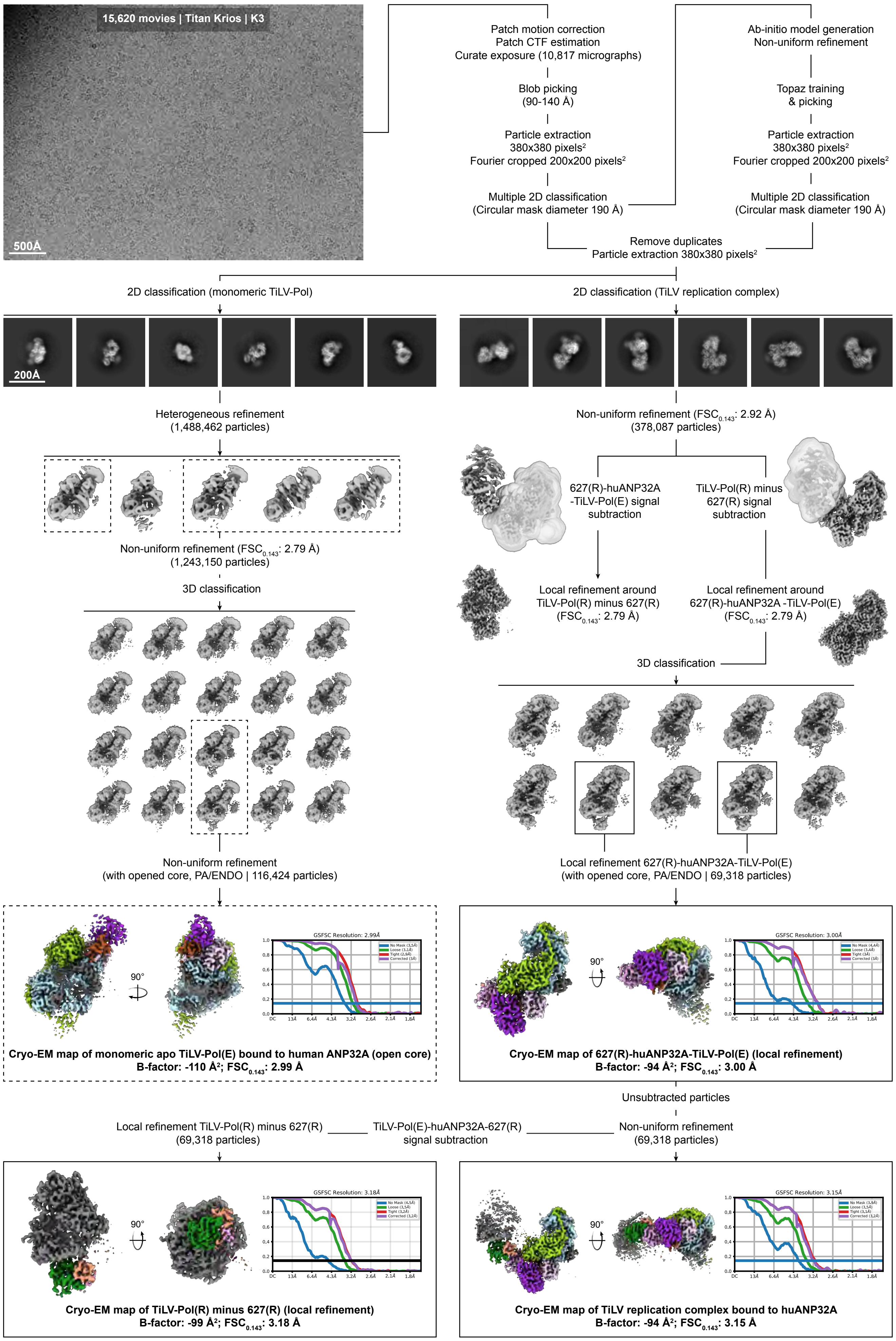

### SUPPLEMENTARY INFORMATION 5

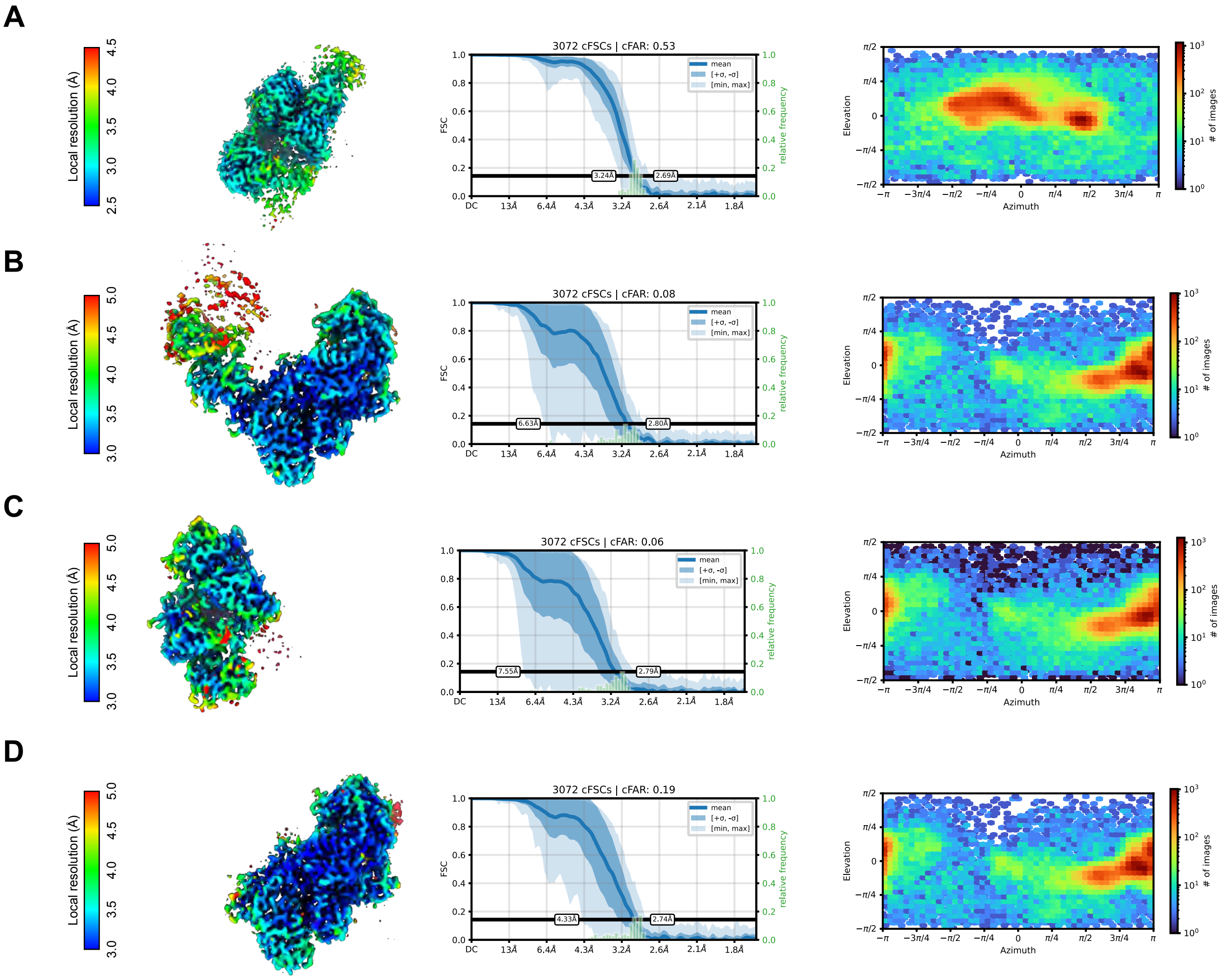

### SUPPLEMENTARY INFORMATION 6

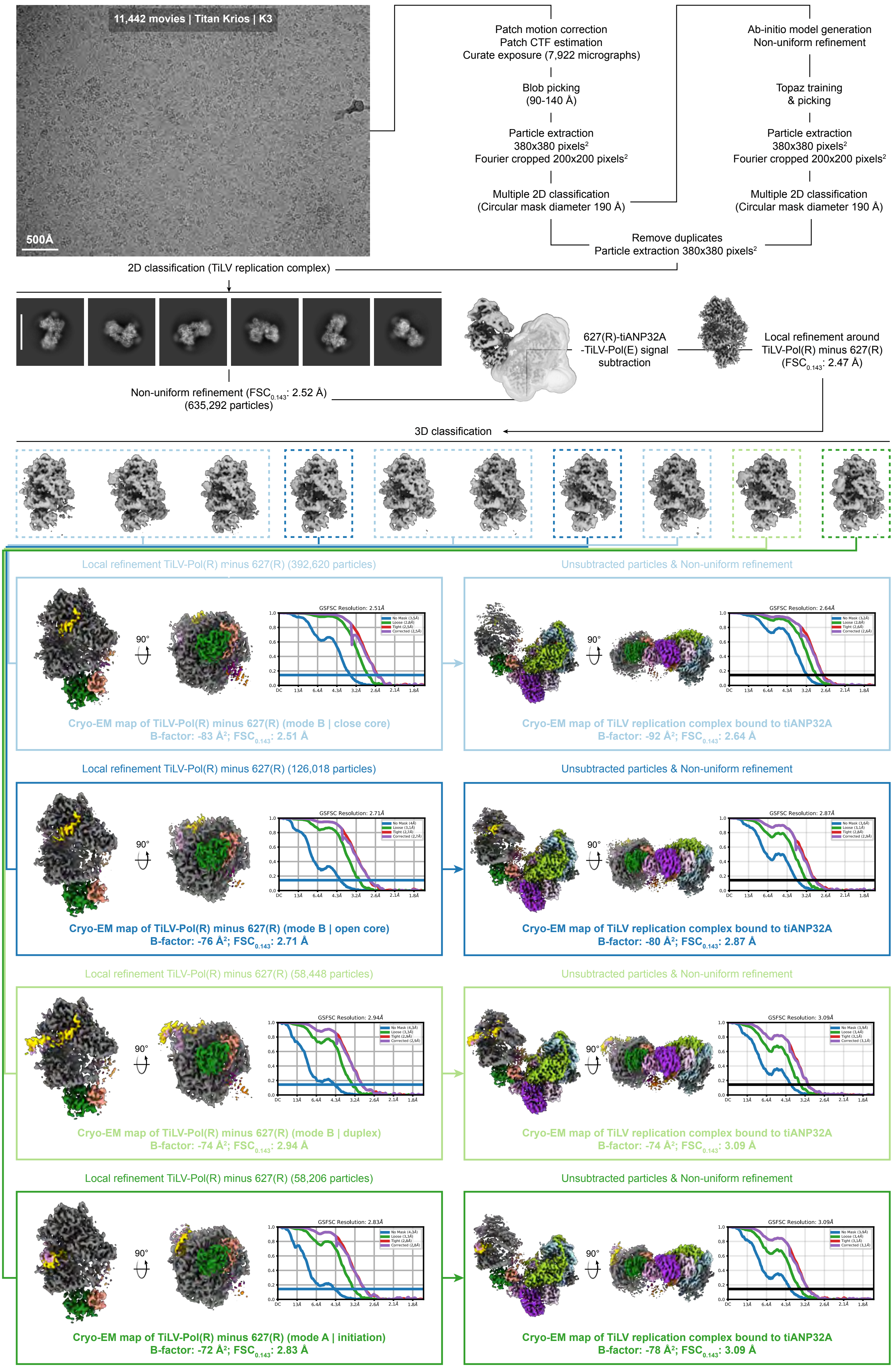

### SUPPLEMENTARY INFORMATION 7

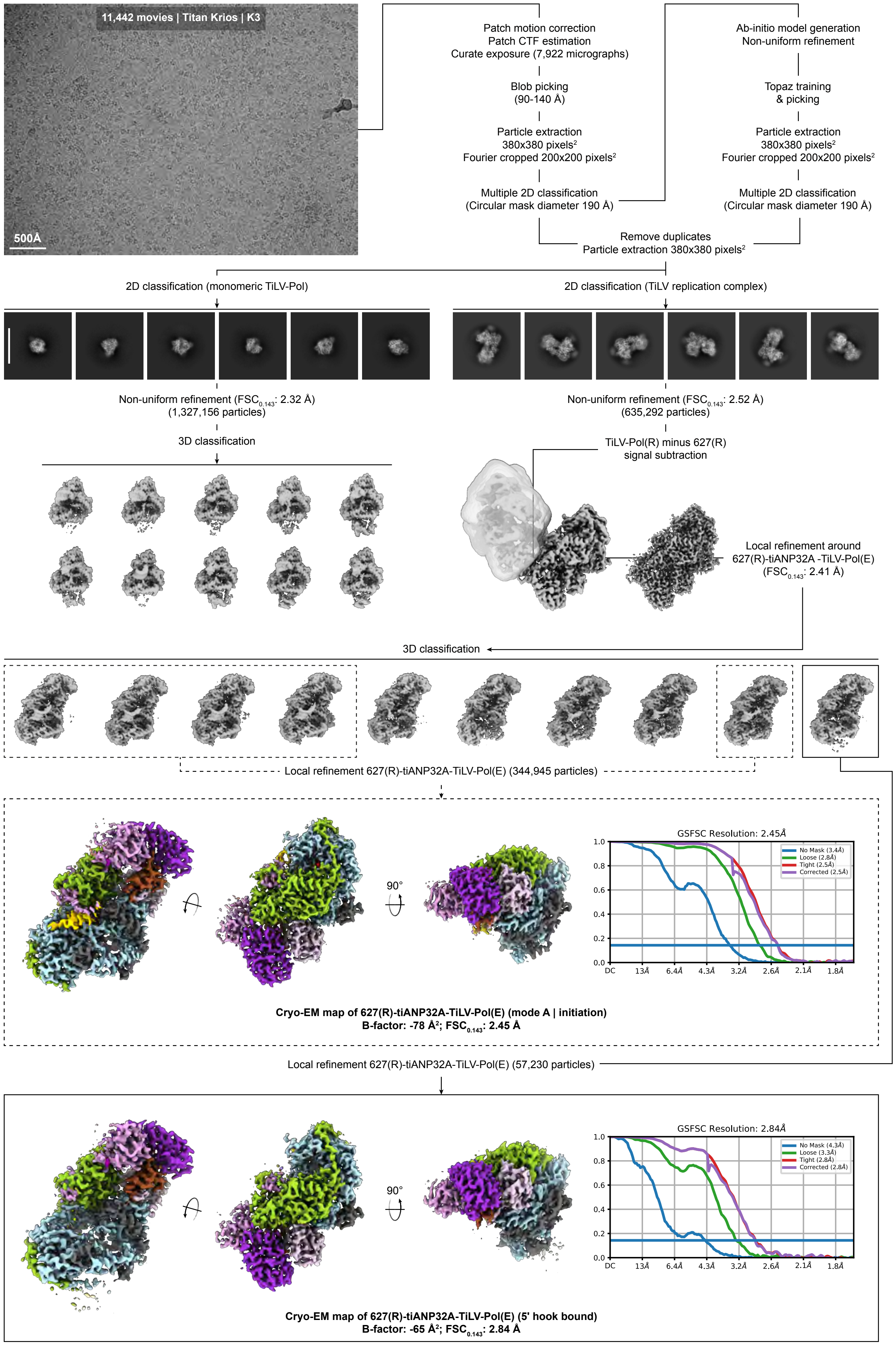

### SUPPLEMENTARY INFORMATION 8

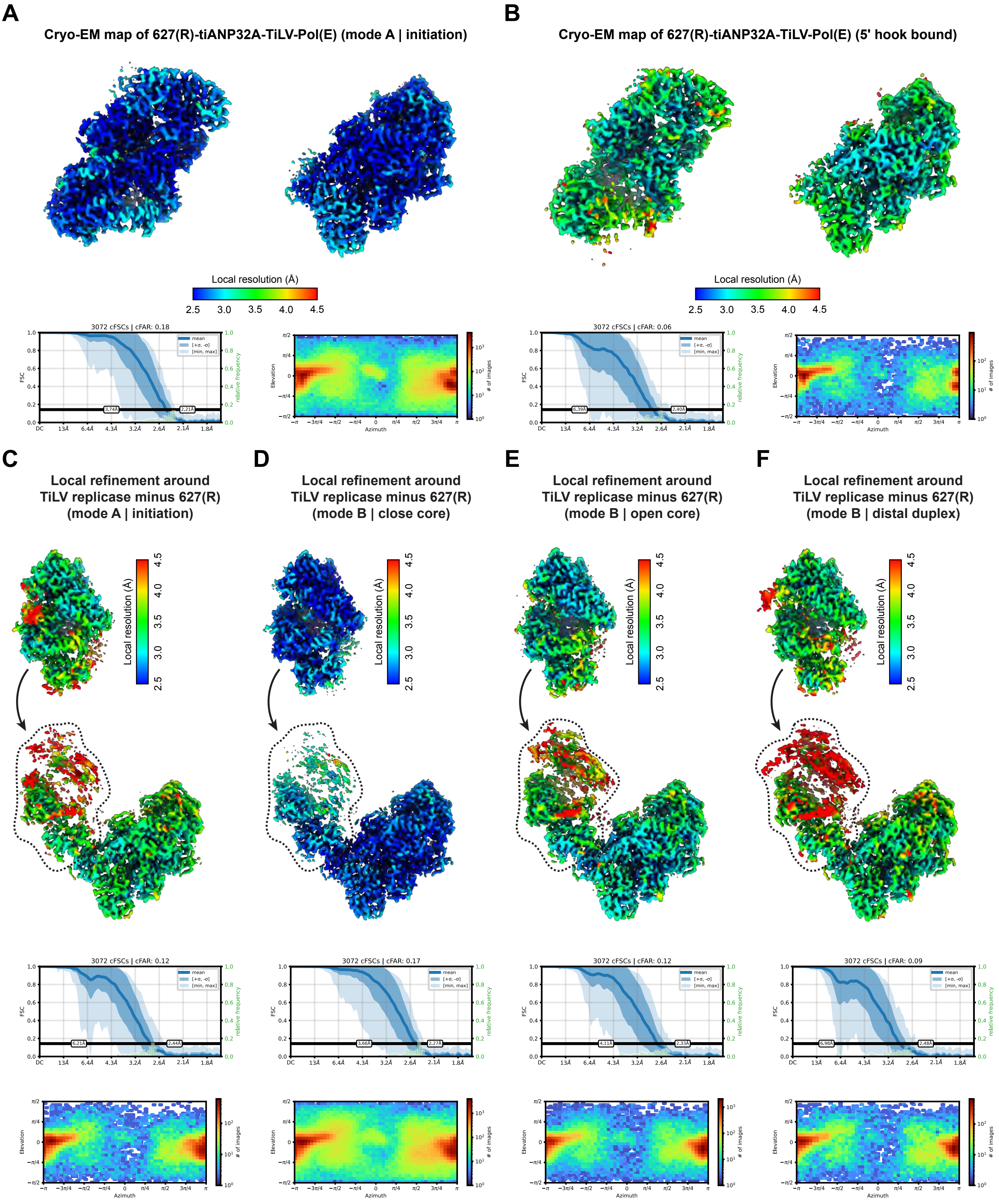

### SUPPLEMENTARY INFORMATION 9

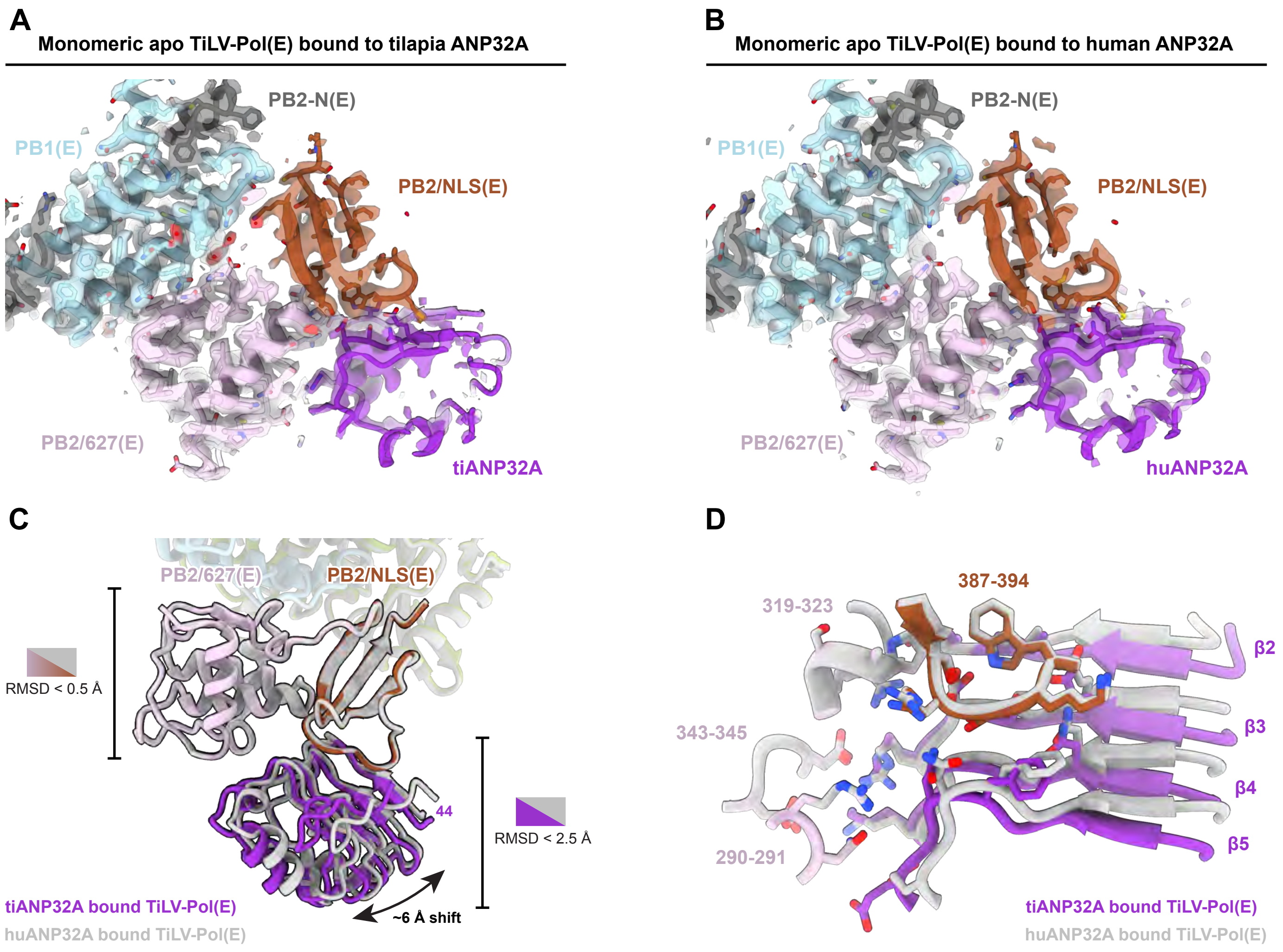

### SUPPLEMENTARY INFORMATION 10

**A**

**Interface between PB2/627-NLS(E) and PA-C(E)**

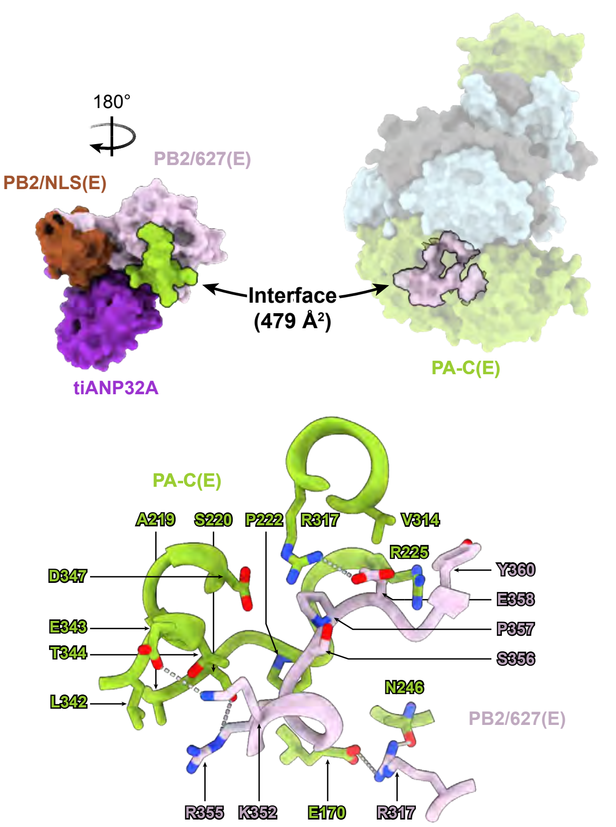


**B**

**Interface between PB2/627-NLS(E) and PB1-C(E)**

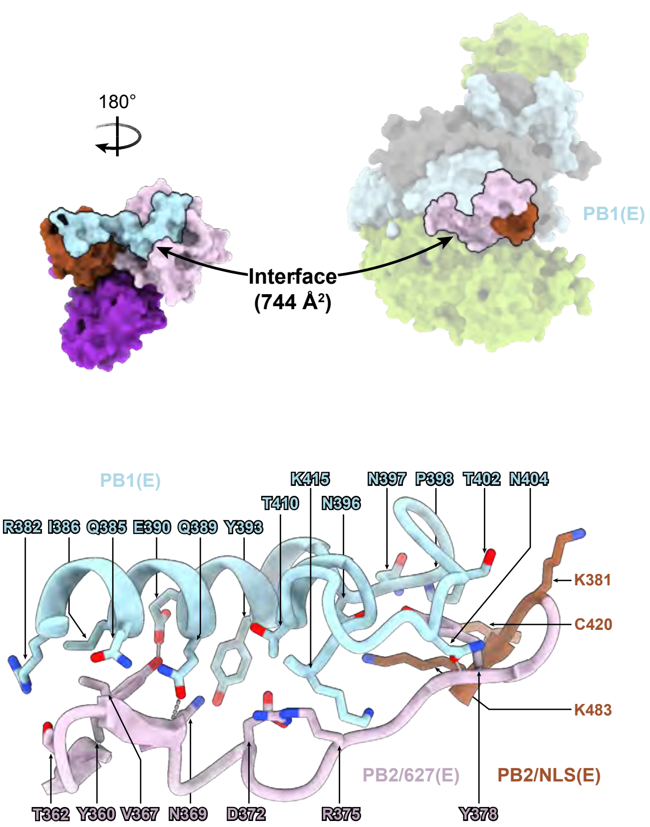


**C**

**TiLV-Pol(T) in pre-termination state (8QZ8)**

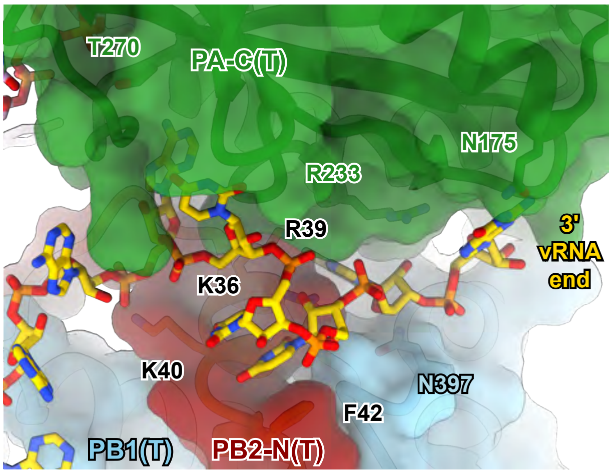


**D**

**Monomeric apo TiLV-Pol(E) bound to tilapia ANP32A**

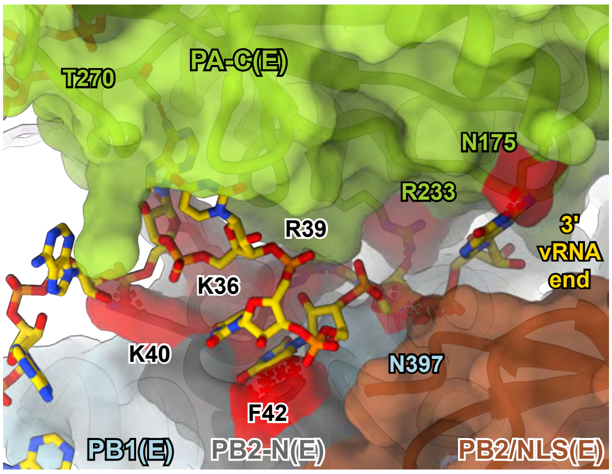

### SUPPLEMENTARY INFORMATION 11

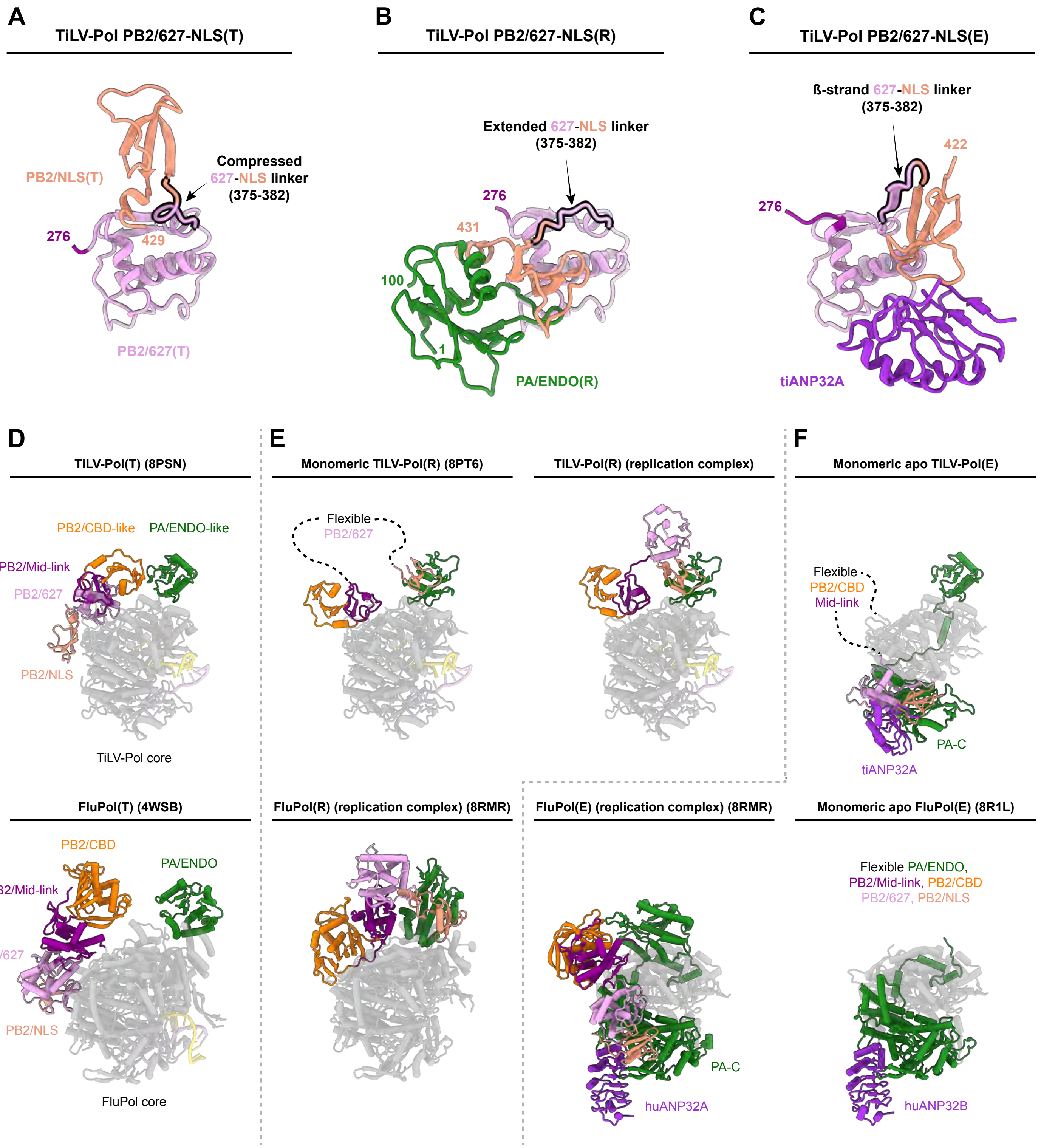

### SUPPLEMENTARY INFORMATION 12

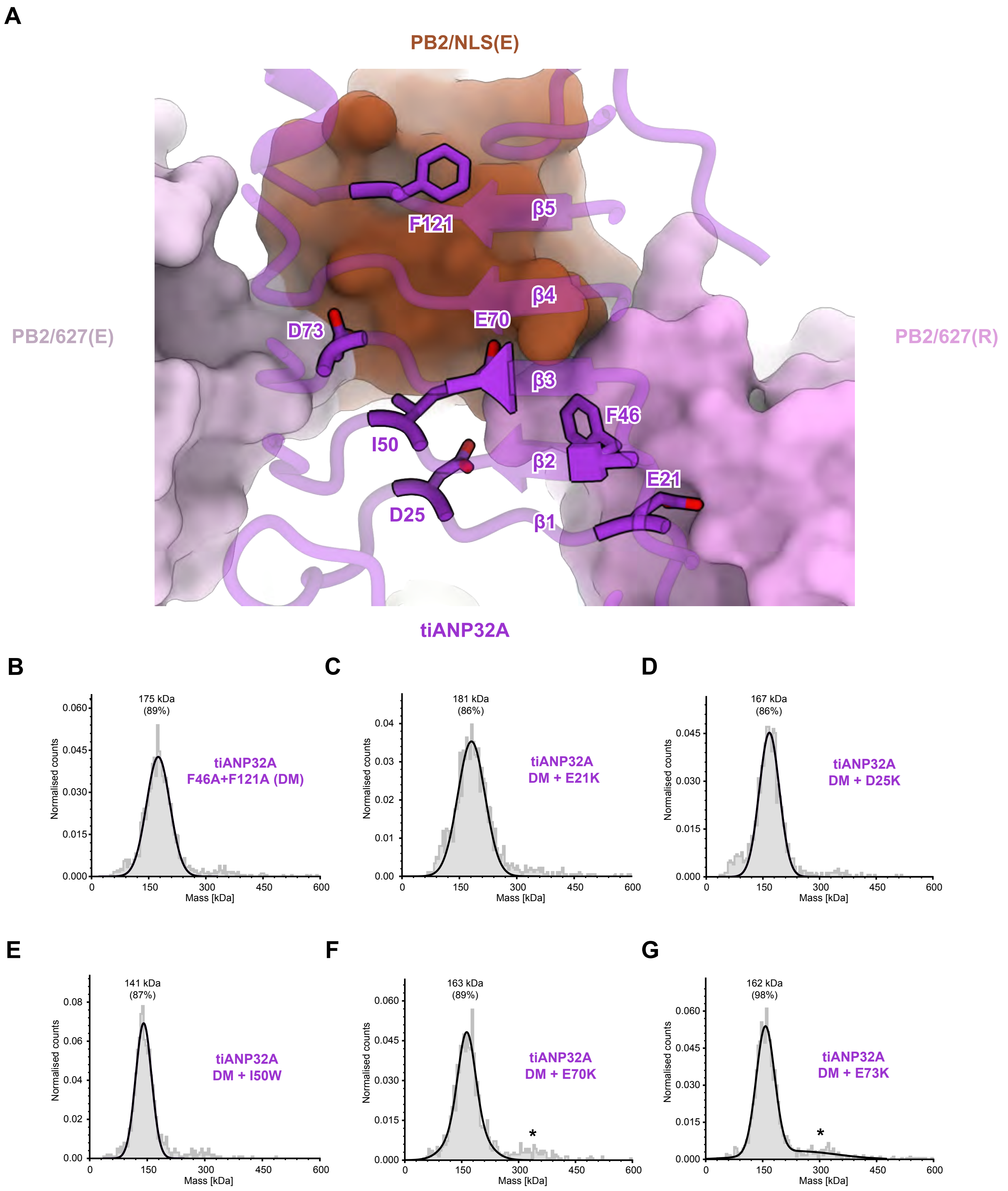

### SUPPLEMENTARY INFORMATION 13

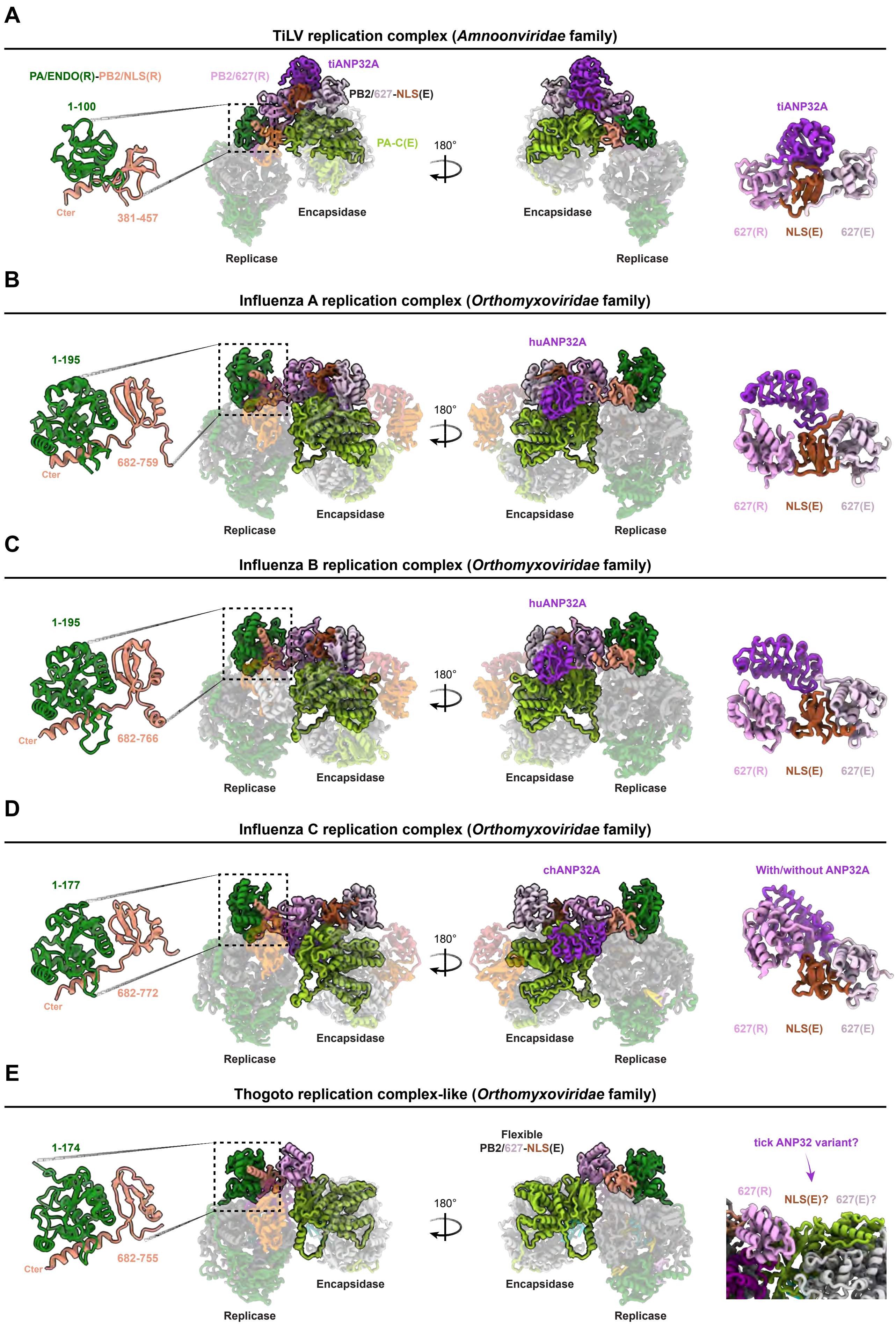

### SUPPLEMENTARY INFORMATION 14

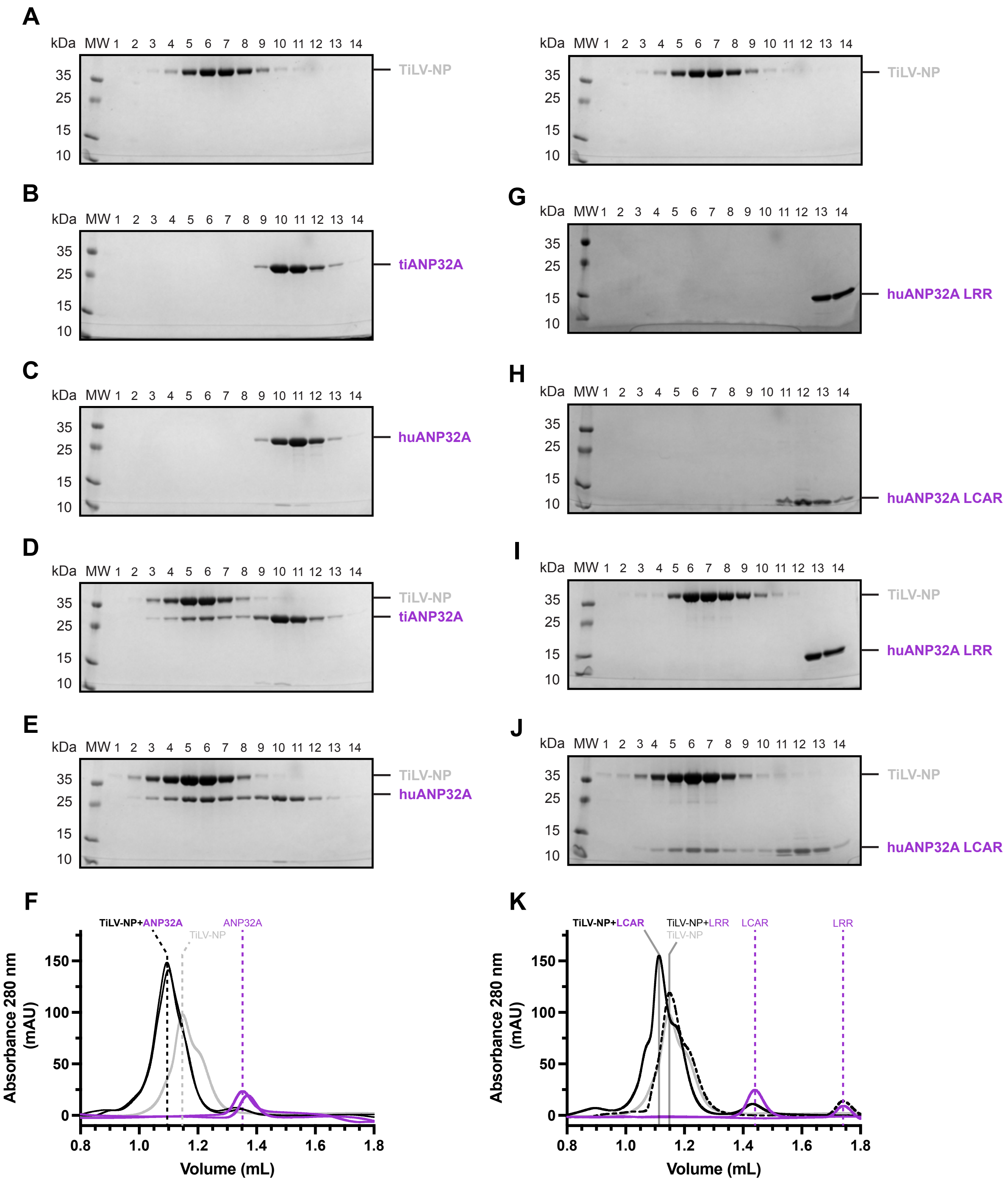
